# Functional Differences in the Neural Substrates of Auditory Cognition as a Consequence of Music Training

**DOI:** 10.1101/2021.06.03.446932

**Authors:** Naomi du Bois, José M. Sanchez Bornot, Dheeraj Rathee, KongFatt Wong-Lin, Mark A. Elliott, Girijesh Prasad

## Abstract

Previous studies have demonstrated that musical deviants (syntactically irregular chords) elicit event related potentials/fields with negative polarity; specifically, the early right anterior negativity and the right anterior temporal negativity responses with peak latencies at ~200 ms and ~350 ms, respectively, post stimulus onset. Here, we investigated differences in the neural dynamics of the auditory perceptual system of individuals with music training compared to those with no music training. Magnetoencephalography was used to examine the neural response to a deviant sound when the auditory system was primed using stimulus entrainment to evoke an auditory gamma-band response between 31 Hz and 39 Hz, in 2 Hz steps. Participants responded to the harmonic relationship between the entrainment stimulus and the subsequent target stimulus. Gamma frequencies carry stimulus information; thus, the paradigm primed the auditory system with a known gamma frequency and evaluated any improvement in the brains response to a deviant stimulus. The entrainment stimuli did not elicit an early right anterior negativity response. Furthermore, the source location of the event-related field difference during the later right anterior temporal negativity response time-window varied depending on group and entrainment condition. In support of previous findings from research using this, and a functionally similar visual-priming paradigm, a 7 Hz phase modulation of gamma amplitude was found for non-musicians following 33 Hz stimulus entrainment. Overall, significant effects of gamma entrainment were found more frequently in the non-music brain. By contrast, musicians demonstrated a greater range of interactions with slower brain rhythms, indicative of increased top-down control.

**Graphical Abstract:** 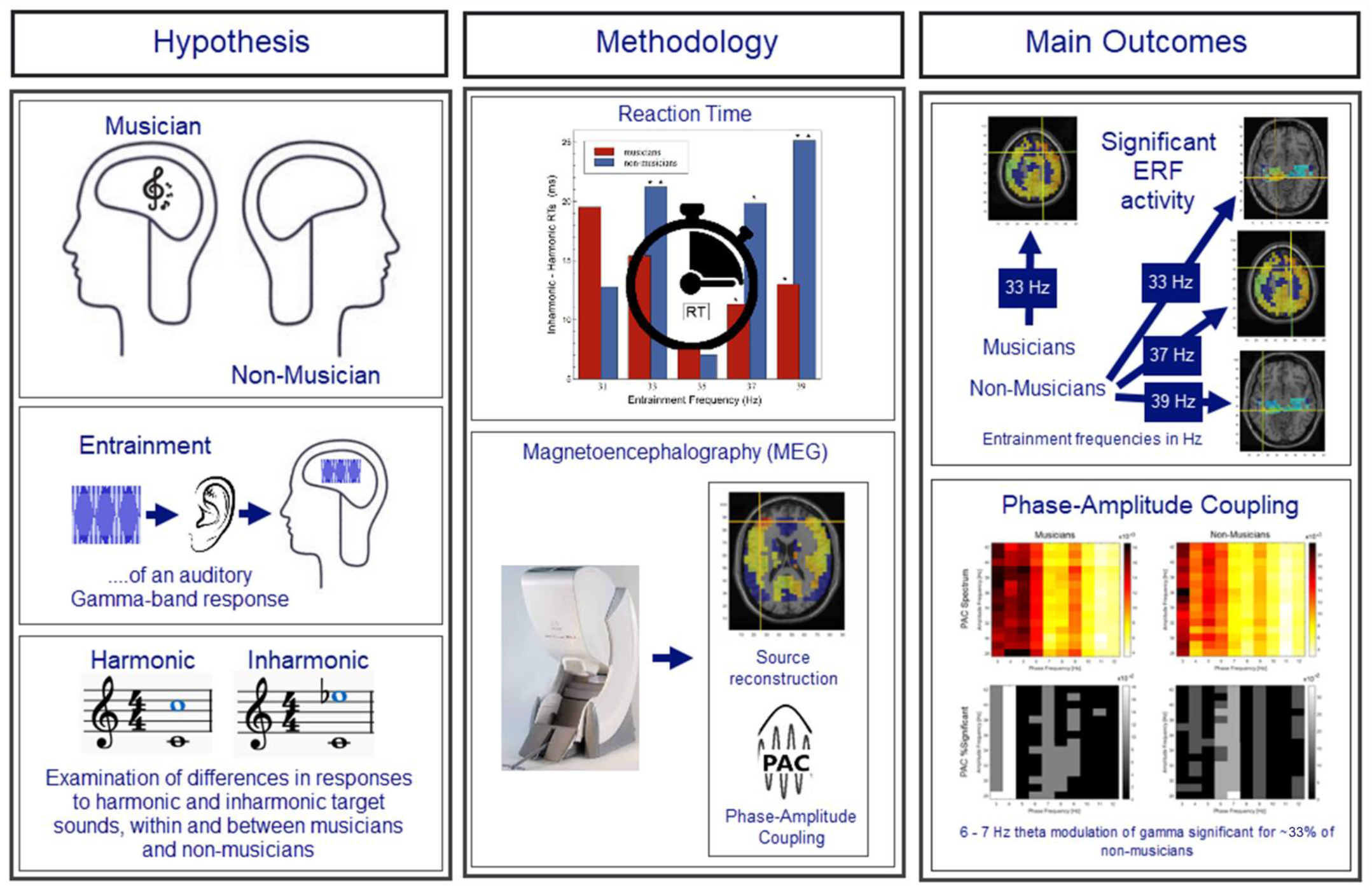

*Conclusion:*

1. Musicians’ auditory cognition relies more on top-down processes, while non-musicians rely on bottom-up processing, and therefore, their auditory cognition is facilitated by the entrained gamma-band response.
2. First neural evidence in support of faster reaction-time responses due to an interaction in phase of an entrained gamma-band response of 33 Hz and a slower endogenous theta rhythm (~7 Hz) - consistently reported in previous research.

## Introduction

Music perception relies on several processes to decode the acoustic information present in the auditory scene such as musical syntax and semantics. These processes include auditory memory and extend to stimulating pre-motor representations of action (Koelsch, 2011). Music cognition is the area of research involved in determining the cortical processes underlying music perception (Justus & Bharucha, 2002). Predictably, musicians, through their training, have developed more efficient cortical mechanisms to subserve these processes. The acquisition of a skill necessarily results in structural and functional changes in the brain to optimise the processes involved in the performance of that skill over time (Pi et al., 2019; Rosenberg-Lee et al., 2018). Regarding harmonic and melodic prediction, neurophysiological music research has demonstrated that syntactically irregular chords elicit event-related potentials (ERPs) with negative polarity and peak latencies of around 150-350 ms post-stimulus onset (Rohrmeier & Koelsch, 2012). The early right anterior negativity (ERAN), and the right anterior temporal negativity (RATN) responses are passively evoked responses, elicited by musical syntax violations. The former arises ~150 ms post-stimulus onset in electroencephalography (EEG) recordings and ~ 200 ms in magnetoencephalography (MEG) recordings, while the latter arises ~ 300 ms post-stimulus onset in EEG recordings and ~ 350 ms in MEG recordings, (Maess, Koelsch, Gunter, & Friederici, 2001; Rohrmeier & Koelsch, 2012). Both the ERAN and RATN response are measured as the difference in cortical activation in response to deviant tones compared to a standard tone. The former is elicited by violations in musical syntax, while the latter occurs when the position of an irregular tone structure within a sequence is unknown, or unpredictable. The RATN response is associated with applying syntactical rules (Koelsch & Mulder, 2002). Given the large body of research documenting structural and functional differences in musicians’ brains compared to their non-musician peers (Angulo-Perkins et al., 2014; Gaser & Schlaug, 2003; Schlaug et al., 1995; Schneider et al., 2005; Zatorre et al., 2007) – a comparison of the music versus the non-music brain provides an excellent means to examine the oscillatory mechanisms involved in auditory binding.

The auditory-priming paradigm used in the current study, to investigate differences in these oscillatory mechanisms as a result of neural plasticity due to music training, was originally designed by Aksentijevic et al. (2011), for the examination of frequency and phase interactions involved in topdown modulation of the oscillatory neural mechanisms that amplify or attenuate incoming stimulus information. The stimuli used in this paradigm are pip-train stimuli, which are effectively short bursts of pure tones that are amplitude modulated, with the amplitude modulation (AM) frequency determined by the duration of the pip, i.e., a pip duration of 25 ms is equivalent to an AM frequency of 40 Hz. Galambos, Makeig, & Talmachoff (1981) discovered when 25 ms pips (which were referred to as ‘clicks’) of different frequencies [250, 350, and 500 Hz] were presented repeatedly to the auditory system, a stable composite 40-Hz event-related potential (ERP) was evoked 8–80 ms post-stimulus onset, that was phase locked to the auditory stimulus which produced the response – termed an auditory (evoked) gamma-band-response (aGBR – see Aksentijevic et al., 2013, 2014; Robert Galambos, 1992). Importantly, a feature of pip-trains is that each pip can carry a different frequency, meaning the carrier frequency (CF) can change with each cycle of the AM frequency. The Aksentijevic et al. (2011) paradigm used these characteristics of pip-train stimuli to establish an entrained aGBR in auditory cortex of a known gamma-band frequency (regulated by the AM of the pip-train), and subsequently examined the frequency-specific effects this had on reaction-time (RT) responses to tones that are either consonant or dissonant with the entrainer pip-trains CFs. Therefore, research using this paradigm refers to the AM frequency as the entrainment frequency. In the paradigm designed by Aksentijevic et al. (2011) two pip-train stimuli are presented separated by a brief (100 ms) inter-stimulus interval (ISI). Both share the same entrainment frequency (in the gamma range). The first pip-train is the entrainer stimulus and carries a repeated four-pip sequence of a single 1000 Hz (deviant) pip followed by three 500 Hz (baseline) pips. The pip-train that follows is the target stimulus, of which there are three types defined by the structure and the harmonic relations between the entrainer and the target – the relation is determined by the interval between the targets deviant CF of 1000 Hz and the entrainers deviant CF. The target absent stimulus contains a single CF of 1000 Hz – thus, there is no interval between this and the entrainer’s deviant CF – the tonal pattern is flat, and the target is absent. A harmonic target has CFs that alternate between a 1000 Hz (baseline) and a 2000 Hz (deviant) – as 2000 Hz is the first harmonic of 1000 Hz, this interval difference is harmonic. An inharmonic target has CFs that alternate between 1000 Hz (baseline) and 2400 Hz (deviant) CFs – as the deviant frequencies in this case are separated by an octave plus a minor third, the interval difference is inharmonic. Both the harmonic and the inharmonic are referred to as target present conditions given the presence of a deviant CF that bears a harmonic relation to the CFs of the entrainment stimuli (see Aksentijevic et al. 2014 for details). Participants are asked to passively listen to the entrainer and then make a speeded response to the target pip train – ‘absent’ if it is monotonic (1000 Hz CF only), or ‘present’ if it has an alternating tonal pattern. The inharmonic violates the listener’s expectancy within a harmonic context, in the same way a ‘deviant’ tone is incongruous compared to a ‘standard’ tone in an oddball scenario. The target absent condition is not of interest. However, it is easier to make a quick and accurate response to the difference in tonal pattern, thus allowing an examination of pre-attentive processing of harmony relations. Stimulus entrainment prepares the auditory system in two ways. Firstly, the entrainment frequency establishes a phase locked auditory GBR and gamma frequencies are known to be involved in carrying feedforward stimulus information and prediction errors (Fries, 2015; Michalareas et al., 2016). Thus, stimulus entrainment provides this information carrier in advance of the target stimulus. Secondly, the CFs of the entrainment stimuli prepare the auditory system to expect a 1000-Hz harmonic relation.

Aksentijevic et al. (2011) found that responses to inharmonic targets were significantly faster than to harmonic targets, only for an entrainment frequency of 33 Hz – termed an ‘inharmonic pop-out’. Moreover, Aksentijevic, Smith and Elliott (2014) found that while musicians respond faster to inharmonic compared to harmonic targets, responses to both target types are faster compared to musically naïve controls - and importantly, the difference between responses to harmonic and inharmonic targets were not significant across a range of entrainment conditions (29 Hz, 31 Hz, 33 Hz, 35 Hz, and 37 Hz). This means there was no evidence of a pop-out effect among musicians. Gamma oscillatory activity has been linked to experience-dependent plasticity during the learning period and once the associative memory has been acquired, performance is no longer influenced by gamma activity (Headley & Weinberger, 2011). Aksentijevic and colleagues (2014) refer to these learned harmonic representations as ‘harmonic templates’ which, once formed, render redundant any reliance on gamma oscillatory binding mechanisms when discriminating harmony relations. Furthermore, the ‘pop-out’ effect among non-musicians has been found to occur following inter-stimulus intervals of ~110 ms and ~260 ms post-stimulus onset. Therefore, Aksentijevic and colleagues (2011) referred to this as a rate- and time-specific reaction time (RT) advantage. Rate in this context describes the number of pips per second (pps) in the entrainment stimulus and thus is interchangeable with frequency. Given these time-specific advantages are separated by approximately 150 ms, which is the duration of one cycle of a 6.7 Hz wave, the inference is that the second time advantage was mediated by a theta rhythm (see Aksentijevic et al., 2011, 2013 for details). Evidence from research investigating the optimal frequency and phase conditions required to process object information in the visual scene suggests that the facilitation effect found when the visual system is similarly primed with specific frequencies in the lower gamma-band range (30 Hz – 70Hz) is also influenced by the phase relation of an endogenous theta rhythm (for details see Elliott, 2014).

The current research examined evidence for a reduced reliance on bottom-up gamma-oscillatory activity due to music training related plasticity effects. Both the ERAN and the RATN response occur during specific time windows post-stimulus onset. However, to date these responses have not been examined under stimulus-entrainment conditions. On this basis, it was hypothesised, given a reduction in reliance on feedforward gamma frequency carriers due to music training, that any effect of stimulus entrainment on the ERAN and RATN responses would be greater for non-musicians than musicians. Just as gamma frequencies have been established as feedforward information carriers, slower brain rhythms (i.e., delta, theta, alpha, and beta) have been found to feedback expected, or predicted, representations (Bubic, Yves von Cramon, & Schubotz, 2010; Engel, Fries, & Singer, 2001; Fries, 2015; Michalareas et al., 2016; Sauseng & Klimesch, 2008). Given previous stimulus entrainment research findings support a gamma-theta interaction that promotes a discriminatory stimulus response, it was further hypothesised that evidence of gamma-theta coupling would be found under entrainment conditions that resulted in improved responses, i.e., a pop-out effect. Thus, the current research aimed to examine the neural mechanisms underlying improved performance in responses to tones that are unexpected, following stimulus entrainment, as suggested by the findings of previous research using this auditory paradigm, and a functionally similar visual paradigm (Aksentijevic et al., 2011, 2014; du Bois & Elliott, 2017; Elliott, 2014).

## Material and methods

Experimental sessions were conducted in the Northern Ireland Functional Brain Mapping (NIFBM) facility, at the Intelligent Systems Research Centre, Magee Campus, Ulster University, Derry, Northern Ireland. The study was carried out in accordance with the Declaration of Helsinki. Approval was obtained for the study from both the National University of Ireland, Galway Research Ethics Committee and the Ulster University, Research Governance department. Informed consent was provided by participants prior to taking part. Experimental sessions lasted between 60 and 80 minutes.

### Participants

Twenty-three participants recruited in Galway and Northern Ireland took part in the study (musician *n* = 11, non-musician *n* = 12). Musicians held Royal Music Academy certificates to a level of grade 3 or above (grade 8+ (x4), grade 6 or 7 (x3), and grade 3 or 4 (x4)), or an international equivalent (*n* = 11, six male, *M* age 32.55 years, *SD* 14.16). All played at least one instrument, and nine played regularly. Non-musicians did not have experience playing a musical instrument (*n* = 12, all male, *M* age 37.44 years, *SD* 9.71). For each participant, sessions began with a 20-trial practice session. Following preprocessing, analysis of the reaction time (RT) data comprised a sample of 11 musicians and 10 non-musicians (due to missing trigger information), while the neuroimaging analysis comprised data from 11 musicians and 9 non-musicians (due to insufficient trials for one other non-musician).

### Task paradigm

A trial sequence began with a 500 ms cue (crosshair), followed by presentation of the entrainer pip-train stimulus for a constant period of 1000 ms, then a constant interstimulus interval (ISI) of 100 ms, and finally the target pip-train. The trial ended following a response to the target, or 2000 ms post-target onset if a response was not made before this time (see Figure 1). Participants were presented with 60 trials per condition which resulted in 1200 trials per participant, (5 x 2(2) x 60 = 1200). Participants were required to respond as rapidly and as accurately as possible, using dedicated response keys to indicate whether the second of the two presented to pip-trains contained alternating tones (i.e., was a target), or was monotonic (a non-target). Target-absent data, i.e., responses to non-targets, were removed from all analyses as the focus of the research was on automatic functional differences between the auditory cognitive processes of musicians and non-musicians when making harmony judgements.

**Figure 1.**
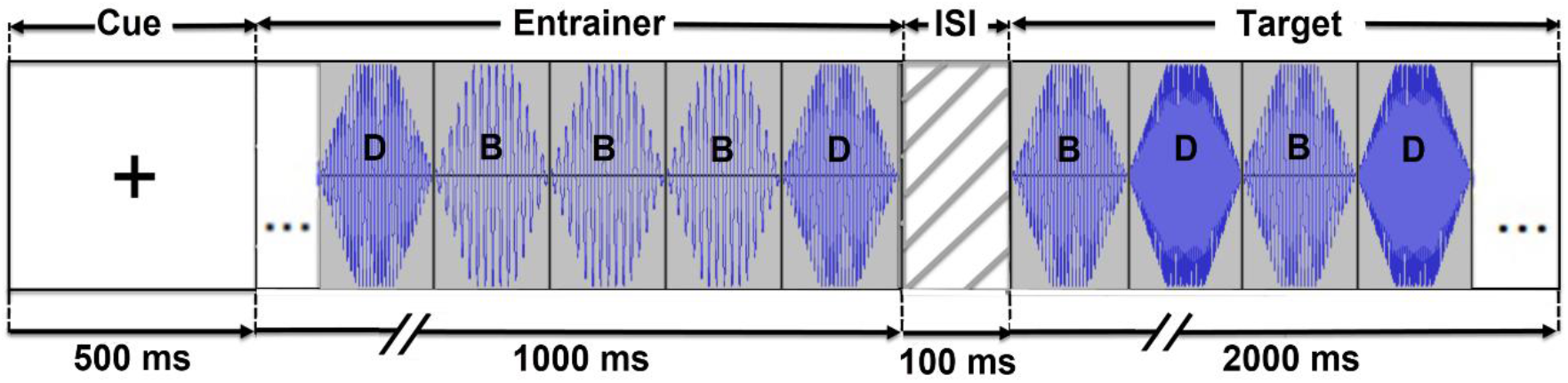
Trial sequence for an inharmonic target in the 33 Hz entrainment condition. The entrainer pip-train for all conditions comprised a repeated pattern of four ‘pips’; one pip with a CF of 1000 Hz (the deviant tone; D) followed by three pips with CFs of 500 Hz (baseline tones; B). The target-absent pip-train for all conditions carried only 1000 Hz pips, while the harmonic- and inharmonic-target pip-trains were composed of an alternating pattern of two pips; the CFs of the alternating pips were 1000 Hz (B) and 2000 Hz (D) in the harmonic target pip-train, and 1000 Hz (B) and 2400 Hz (D) in the inharmonic target pip-train. The amplitude modulation frequency in the above example was 33 Hz – therefore, the 1000 ms entrainment period ended following a deviant pip in the repeated pattern of four pips (DBBB).

The parameters of the auditory pip-train stimuli were designed to replicate the stimuli in the paradigm which produced the previous ‘pop-out’ effect following stimulus entrainment of 33 Hz. Thus, the entrainment frequency range for the current experiment was 31 to 39 Hz in 2 Hz steps. Experimental trial plus stimulus and trigger generation were coded in-house using COGENT 2000 and COGENT GRAPHICS (http://www.vislab.ucl.ac.uk/cogent.php) toolboxes running in MATLAB R2016b (Oostenveld, Fries, Maris, & Schoffelen, 2011). The stimulation PC was a HP xw4600 Workstation.

### Trials with incorrect responses

The percentage number of trials that were incorrectly responded to, was 8.21% overall. The arcsine of the square root of the proportion of incorrect responses for each condition were analysed showing no significant differences in the production of incorrect responses across participants.

### MEG data acquisition

The continuous raw MEG data was recorded per participant, per block (200 trials) using the superconducting quantum interference device (SQUID) based 306-channel whole head MEG Elekta Neuromag TRIUX system (Helsinki, Finland), comprising 204 gradiometers and 102 magnetometers. Head shape was obtained by using a three-dimensional Fastrak digitiser (Polhemus) by acquiring three fiducial (or cardinal) points (nasion and left and right preauricular points) and at least 300 points of the surface of the scalp. In addition, four head position indication (HPI) coils were placed on the subjects’ head: two on the mastoids and two on the forehead. The positions of the HPI coils were acquired using the Fastrak device to provide continuous head position estimation during the recording. Ocular movements and cardiac activity were measured for cleaning purposes using four electro-oculography (EOG) electrodes (horizontal, and vertical pairs), and two cardiac muscle electrodes. Signals were digitised with a bandwidth of 0.1 Hz to 300 Hz and a sampling rate of 1000 Hz. Sound stimuli were presented binaurally via ER-3A ABR Insert Earphones, and the decibel level was attenuated to 50 SPL, as measured by a Precision Gold (IEC 651 TYPE II) sound level meter (model #: N05CC). Instructions and a cross to cue the start of each trial were rear-projected onto a 112 cm screen with a resolution of 1024 9 768 pixels at 60 Hz, using a zero-jitter 3-DLP video projector and lens (RM-MSX21G; Victor Company of Japan, Kanagawa, Japan). The screen was placed approximately 120 cm from the participant, who was seated in the upright position. A bilateral finger response system was used to collect participants RT responses.

Data cleaning and analyses (other than the phase-amplitude coupling (PAC) analysis) were performed using in-house scripts in the Fieldtrip toolbox for Matlab R2016b (Oostenveld et al., 2011). The PAC analysis was performed using the Spatial Parametric Mapping (SPM12) toolbox (Litvak et al., 2011).

### Preprocessing

The continuous raw data were filtered off-line using Max Filter temporal signal space separation (tSSS). Data for each participant were then epoched and preprocessed, removing target-absent and response error trials and offsetting the samples to account for the delay in sound stimuli presentation. A bandpass filter was used to filter out frequencies above 150 Hz, and a discrete Fourier transform was used to filter out line noise. Separate blocks for each participant were concatenated and resampled from 1000 to 500 samples per second. Trials were manually inspected to remove noisy segments, and the data were checked for trials and channels that were outliers. Squid jump detection was used to identify jumpy channels (a cut-off *z* score of 25 was used). Channels that were outliers (due to noise) were marked as bad and the remaining bad trials detected were removed. Muscle artifacts were removed (using a cut-off *z* score of 12), and a final visual inspection was performed. Finally, an independent component analysis (ICA) was performed to identify and remove electro-ocular and electro-cardiac components. The clean data were interpolated to repair channels marked as bad. Data were then averaged across trial conditions, i.e., inharmonic-target present (ITP) by entrainment frequency and harmonic-target present (HTP) by entrainment frequency, for each participant.

### Source analysis

A source reconstruction analysis was performed using a linearly constrained minimum variance (LCMV) beamformer (Van Veen, Van Drongelen, Yuchtman, & Suzuki, 1997). In the absence of a T1 MRI for the participants, a 1 mm resolution template MRI was used to create the head model and 3D source model. The template MRI was co-registered to the MEG coordinates using the three fiducial points (i.e., nasion, left preauricular, and right preauricular point). The registered images were normalised to the MNI template using a nonlinear registration algorithm (Ashburner, 2007) and then used to define a homogeneous grid of 2 mm resolution. A single shell volume conduction model was created based on the segmentation of the head tissues. Lead fields were defined using the single shell head model to fit the head shape of each subject in the vicinity of each sensor. Spatial filter coefficients were estimated for each participant using the computed lead field and an average of the covariance matrix for all the epochs (both harmonic and inharmonic). Thereafter, this filter was used to compute the sources for each harmonic and inharmonic condition separately. This method results in independent scaling factors for all three spatial dimensions, thus enabling source localization with an accuracy close to that achieved with individual MR-based models. The source activity elicited by harmonic and inharmonic targets, in the time windows corresponding with the ERAN and RATN responses, were averaged over trials within each condition and the Monte Carlo Method was used to calculate the significance probability of variability using the dependent samples t-statistic. The following three analyses were performed in this way, on the source data in both the ERAN and the RATN time windows.

1. To examine inharmonic-harmonic event-related field (ERF) differences overall: Analysis of variance was performed between the source activations elicited by the inharmonic compared to the harmonic, at each entrainment frequency for the combined sample.
2. To examine inharmonic-harmonic ERF differences within each group separately: Analysis of variance was performed between the source activations elicited by the inharmonic compared to the harmonic targets for each entrainment frequency in each group separately.
3. To examine inharmonic-harmonic ERF differences between musicians and non-musicians: Analysis of variance was performed between the inharmonic-harmonic source activity differences, at each entrainment frequency, dependent on music group.

The design matrix for each analysis was for a within-subject manipulation. In order to have equal units of observation for the musicians versus non-musicians matrix (analysis number three), two musicians (participants 3 and 9) were randomly selected, in Matlab, to be excluded from the musician sample.

The source analysis is focused on inharmonic-harmonic differences for each entrainment frequency, dependent on music group, i.e., within-group effects investigating inharmonic-harmonic source activity differences for each group, and between group effects examining group differences in the ERF difference activity. Furthermore, all analyses were performed over the time windows associated with both the early ERAN response (~200 ms) and the later RATN (~350 ms) response. The methodology is designed to assess functional (strength of a response) and structural (location of a response) differences in the source activations in response to an inharmonic stimulus that is dependent on music training. The results of analyses conducted on data within the ERAN time window (t = [175 –225 ms]) did not reach significance at an alpha level of .05.

### Phase-amplitude coupling analysis

For the PAC analysis the sources were computed for both harmonic and inharmonic trials combined, and across a time window of −1000 to 500 ms. This source data was parcellated and used to generate virtual sensors. General Linear Models (GLMs) were estimated for centre frequencies between 3 and 12 Hz, in steps of 1 Hz, for the phase-frequency component and between 28 and 42 Hz with 1 Hz steps for the amplitude-frequency component, with a frequency resolution of .5 Hz. All epochs were concatenated to estimate the overall cross-frequency coupling, while the single epochs were entered individually into a separate GLM to test for significance at two alpha levels: .01, and .05. PAC correlation coefficients (r_pac_) were computed for participants in each (entrainment) condition (van Wijk, Jha, Penny, & Litvak, 2015). Group averaged comodulograms were generated, both by averaging across the individual r_pac_ images, and by averaging across the individual sig_pac_ images. A series of paired t-tests were performed on the r_pac_ images to examine statistical differences between conditions within each group (musicians and non-musicians) (for details see, van Wijk et al., 2016)

## Results

### Reaction Time results

Examination of the raw RT data using the Statistical Package for Social Sciences (IBM SPSS statistics 23.0) revealed a non-normal distribution with a positive skew for data in the 35 Hz condition (*W*(20) = .889, *p* = .026, the absolute skewness value for the data in this condition was .467). Subsequent analyses were conducted on the exponents of the means of log-transformed RT distributions, calculated in MATLAB using an in-house script (for supporting ideas see Box, & Cox, 1982; Box & Cox, 1964). A mixed repeated-measures analysis of variance (ANOVA) was applied to these data to evaluate the effect of entrainment frequency (31 Hz, 33 Hz, 35 Hz, 37 Hz, and 39 Hz), and harmonic relation (harmonic and inharmonic targets) on RT responses, as a function of music experience (group being the between factor; musicians and non-musicians). Levene’s test for equality of variance was applied to the analysis, as was Greenhouse-Geisser estimates of sphericity, and degrees of freedom were corrected when a violation to an assumption was indicated. Bonferroni correction was applied to all pairwise comparisons.

Significant main effects were found for target, and entrainment frequency; *F*_(1, 19)_ = 6.13 *p* = .023, *η_p_^2^* = .24, *F*_(4, 76)_ = 2.79, *p* = .032, *η_p_^2^* = .13 respectively. The effect of group was not found to be significant (*F*_(1, 19)_ = .004, *p* = .95). The main effect for target was due to slower responses to harmonic compared to inharmonic targets (*MD* = 15.32, *SE* = 6.19, *p* = .023). The main effect for entrainment frequency was due to faster RTs to targets with an entrainment frequency of 35 Hz compared to 31 Hz (*MD* = 11.9, *SE* = 3.25, *p* = .016). While the interaction between target and entrainment frequency was not significant (*p* = .38), a pairwise comparison revealed significant RT differences between targets following 33 Hz, 37 Hz, and 39 Hz entrainment (*MD* = 18.31, *SE* = 6.68, *p* = .013, *MD* = 15.58, *SE* = 7.14, *p* = .042, and *MD* = 19.08, *SE* = 7.59, *p* = .021, respectively). While the three-way interaction between group, target and entrainment frequency did not achieve significance (*p* = .57), a pairwise comparison found the target by entrainment frequency effect (found for 33 Hz, 37 Hz, and 39 Hz entrainment conditions) held following 33 Hz and 39 Hz entrainment – for non-musicians only (*MD* = 21.22, *SE* = 9.66, *p* = .041, and *MD* = 25.14, *SE* = 10.99, *p* = .034, respectively).

As anticipated the results revealed a 33 Hz inharmonic pop-out in the non-musician group data, consistent with that reported by Aksentijevic et al., (2011, 2014). A second pop-out was also found at 39 Hz for non-musicians. Following 35 Hz entrainment the difference in responses for both groups to both targets was very small, with a main effect found which indicated responses for conditions with an entrainment frequency of 35 Hz were significantly faster compared to an entrainment frequency of 31 Hz. Aksentijevic et al., (2014) found a main effect of entrainment frequency for 33 Hz and 35 Hz conditions, thought to reflect attentional mechanisms which rely on the temporal features of the sound, communicated by the envelope fluctuations of the entrainment (AM) frequency. Furthermore, the musicians’ responses were not faster overall compared to non-musicians, contrary to the responses of the musicians in the Aksentijevic et al., (2014) sample of jazz musicians. However, while jazz and classical musicians have been found to possess similar levels of tonal ability, jazz musicians demonstrate a greater sensitivity to auditory deviants - resulting in a larger auditory mismatch-negativity (MMN) response (Vuust, Brattico, Seppänen, Näätänen, & Tervaniemi, 2012). It is considered this could result in greater accuracy and faster responses to the auditory stimuli used within the paradigm, although further investigation would be needed to clarify.

The main finding from the analysis of the RT data is that entrainment of 33 Hz, 37 Hz, and 39 Hz resulted in responses to inharmonics that were significantly faster compared to harmonic responses. Furthermore, a group comparison showed that musicians’ responses to harmonics and inharmonics did not differ significantly across all entrainment conditions, while non-musicians demonstrated faster responses to inharmonic than to harmonic target tones, following 33 Hz and 39 Hz entrainment.

### Source analysis

Source locations from MEG activity, found to be significant during the RATN time window (t = [325 – 375 ms]), were inconsistent with the expected RATN location, i.e., right frontotemporal region as described by Patel et al. (1998). However, significant inharmonic-harmonic ERF differences were found within the RATN time window during entrainment conditions that were consistent with significant inharmonic-harmonic RT outcomes. To recap, analysis of the RT data revealed faster responses to the inharmonic targets following entrainment at 33 Hz, 37 Hz, and 39 Hz entrainment frequencies. However, a multiple comparisons analysis of the factor-level effects determined a priming effect due to significantly faster responses to inharmonic compared to harmonic targets following 33 Hz and 39 Hz entrainment, for non-musicians only.

Firstly, the analysis of variance between the ERF generated by the inharmonic compared to the harmonic, for the combined sample (musicians and non-musicians), found the strength of the response to be increased for the inharmonic following entrainment frequencies of 33 Hz, 37 Hz and 39 Hz (33 Hz, cluster statistic = - 3.8905 x10^3^, *p* = .015; 37 Hz, cluster statistic = 1.3703 × 10^3^, *p* = .0043; 39 Hz, cluster statistic = - 973.086, *p* = .044). This finding was replicated in the second analysis of variance among the non-musician cohort only, following entrainment frequencies of 33 Hz and 39 Hz (33 Hz, cluster statistic = - 2.749 ×10^3^, *p* = .022; 39 Hz, cluster statistic = - 878.7248 ×10^3^, *p* = .037).

When a threshold alpha-value of 0.05 was adopted for the dependent samples t-statistic, to initially threshold the data for the cluster analysis, a group difference did not reach significance. However, inharmonic-harmonic ERF source differences that have similar activation strengths and locations for both groups will go undetected, even though the ERF difference may be significant for both groups, simply as they will not be significantly different for one group compared to the other. Lowering the initial thresholding alpha value to 0.01 to restrict the critical values entered in the cluster-based permutation routines, allows for more subtle group differences to be detected (for details see Gaab & Schlaug, 2003). Taking this approach, significant group differences were found for both groups - in separate locations, following 33 Hz entrainment. A significant variance in inharmonic-harmonic ERF source activity was found for musicians compared to non-musicians, in the middle frontal gyrus (cluster statistic = 221.018, *p* = .033, see Figure 5), and for non-musicians compared to musicians in the left parahippocampal and left fusiform gyri (cluster statistic = −221.018, *p* = .05, see Figure 6).

**Figure 2.**
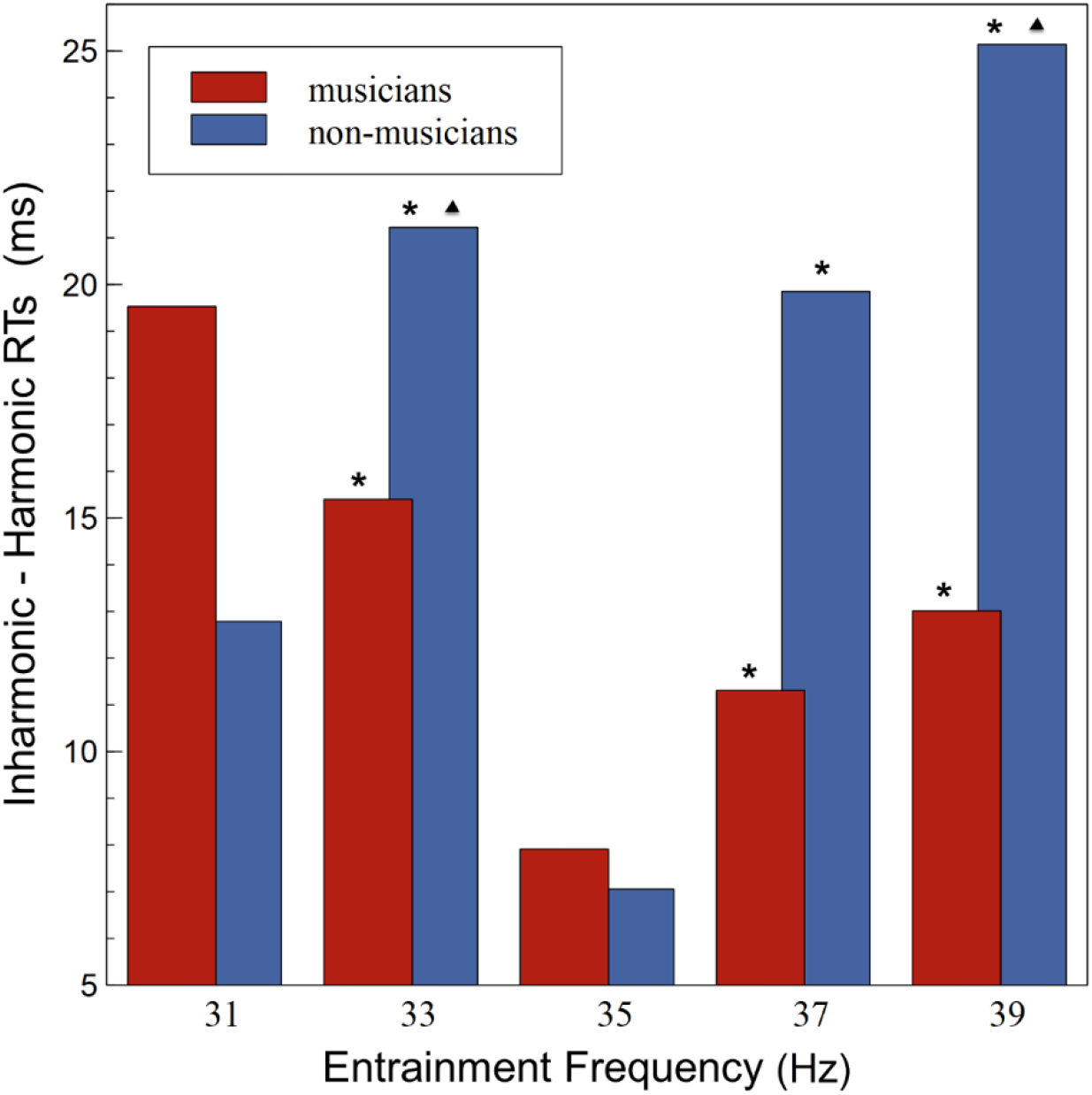
Differences in average reaction times (RTs) between harmonic and inharmonic responses, for each entrainment condition and group. Averaged harmonic RT responses subtracted from averaged inharmonic RT responses for musicians (red bar) and non-musicians (blue bar) for entrainment frequency condition (x-axis). Pairwise comparison found faster responses to inharmonics following 33 Hz, 37 Hz, and 39 Hz entrainment (*), while pairwise comparison of all three factors (group, target and entrainment frequency) revealed the RT difference remained significant for non-musicians only following 33 Hz and 39 Hz entrainment (triangles).

**Figure 3.**
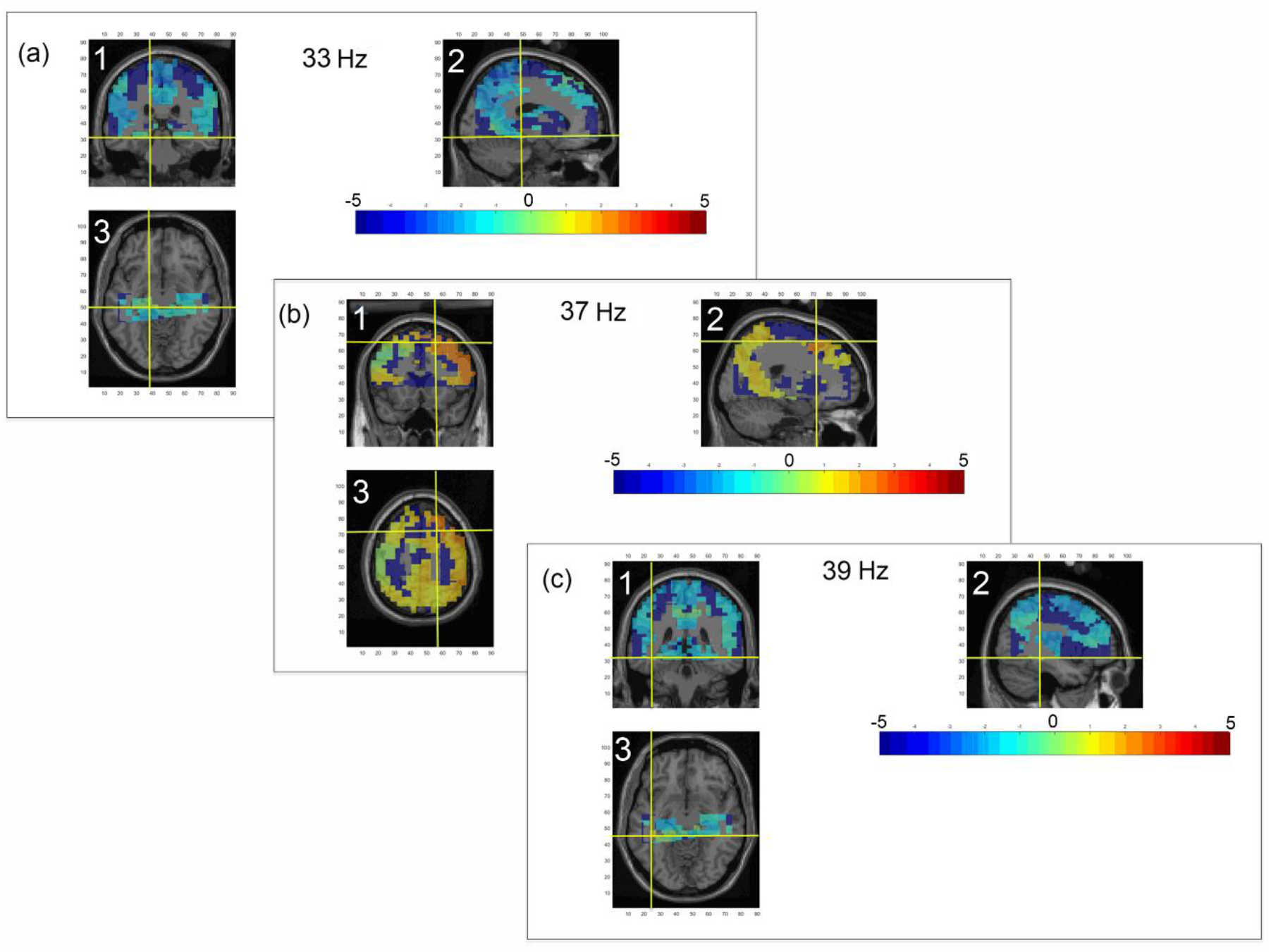
Mean images depicting the MEG source localisation of the difference between harmonic and inharmonic ERF activation, ~350 ms post-stimulus onset (time window = [325 – 375 ms]), for musicians and non-musicians combined, following entrainment frequencies of (a) 33 Hz, (b) 37 Hz, and (c) 39 Hz. For each condition, the source location is presented in (1) coronal (2) sagittal and (3) transverse views. Source activities identified: (a) left hippocampus and parahippocampal area (MNI coordinates [−16 −30 −12] mm), following 33 Hz entrainment; (b) right superior frontal cortex (MNI coordinates [20 16 56] mm), following 37 Hz entrainment; and (c) left inferior temporal lobe (MNI coordinates [− 42 − 38 − 12] mm), following 39 Hz entrainment. Values plotted are the cluster t-statistics within the range −5 to 5 (see colour bar), with significant clusters (*p* < .05) marked by the crosshair.

**Figure 4.**
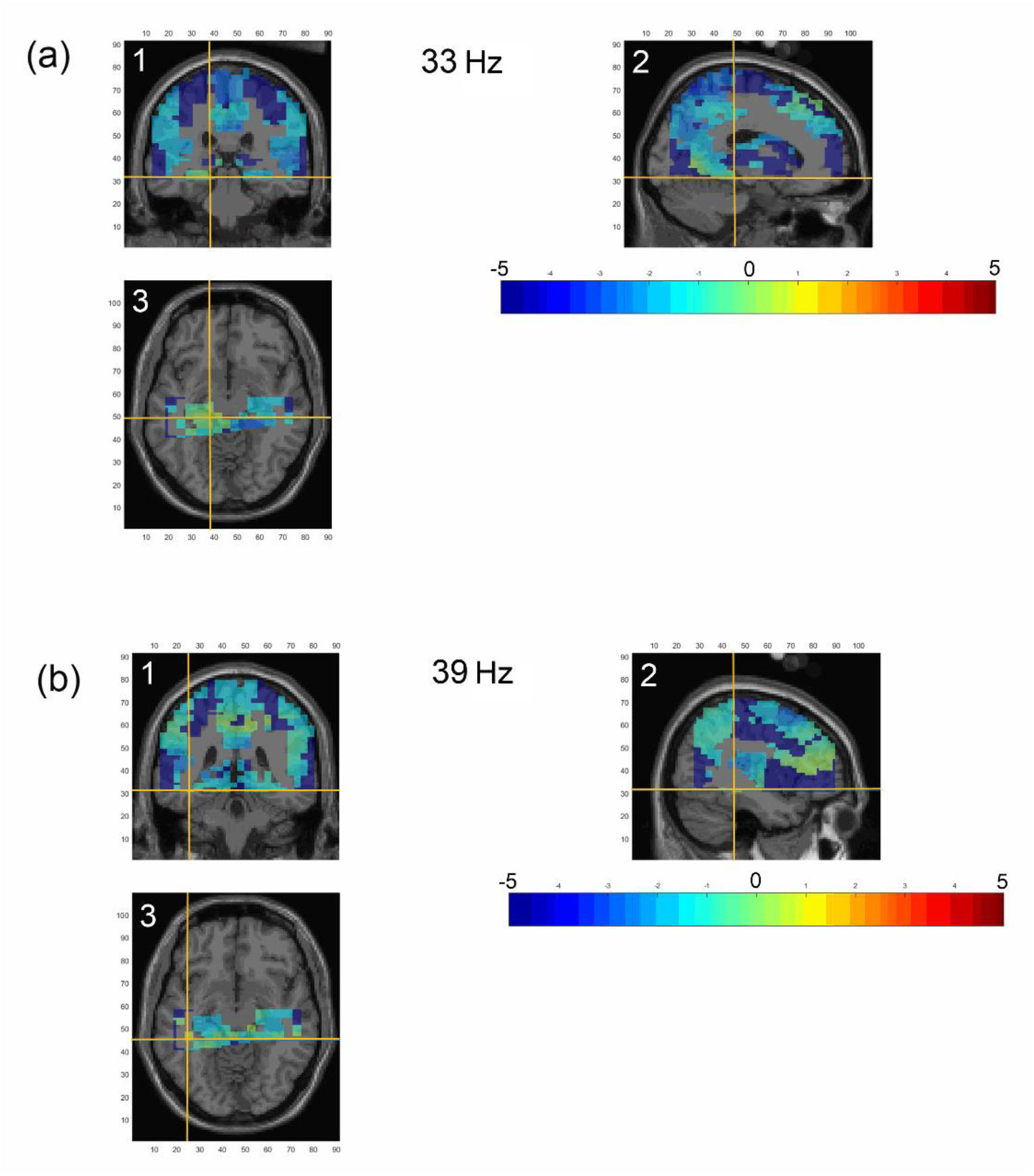
Mean images depicting the source localisation of the difference between harmonic and inharmonic ERF activation, ~350 ms post-stimulus onset (time window = [325 – 375 ms]), following entrainment frequencies of (a) 33 Hz and (b) 39 Hz, for the non-musician group. For each condition, the source location is presented in (1) coronal (2) sagittal and (3) transverse views. As for the combined data under the same entrainment conditions, (a, 33 Hz) is located in the left hippocampus and parahippocampal area (MNI coordinates [−16 −30 −12] mm), and (b, 39 Hz) in the left inferior temporal lobe (MNI coordinates [−42 − 38 − 12] mm). Values plotted are the cluster t-statistics within the range −5 to 5 (see colour bar), with significant clusters (*p* < .05) marked by the crosshair.

**Figure 5.**
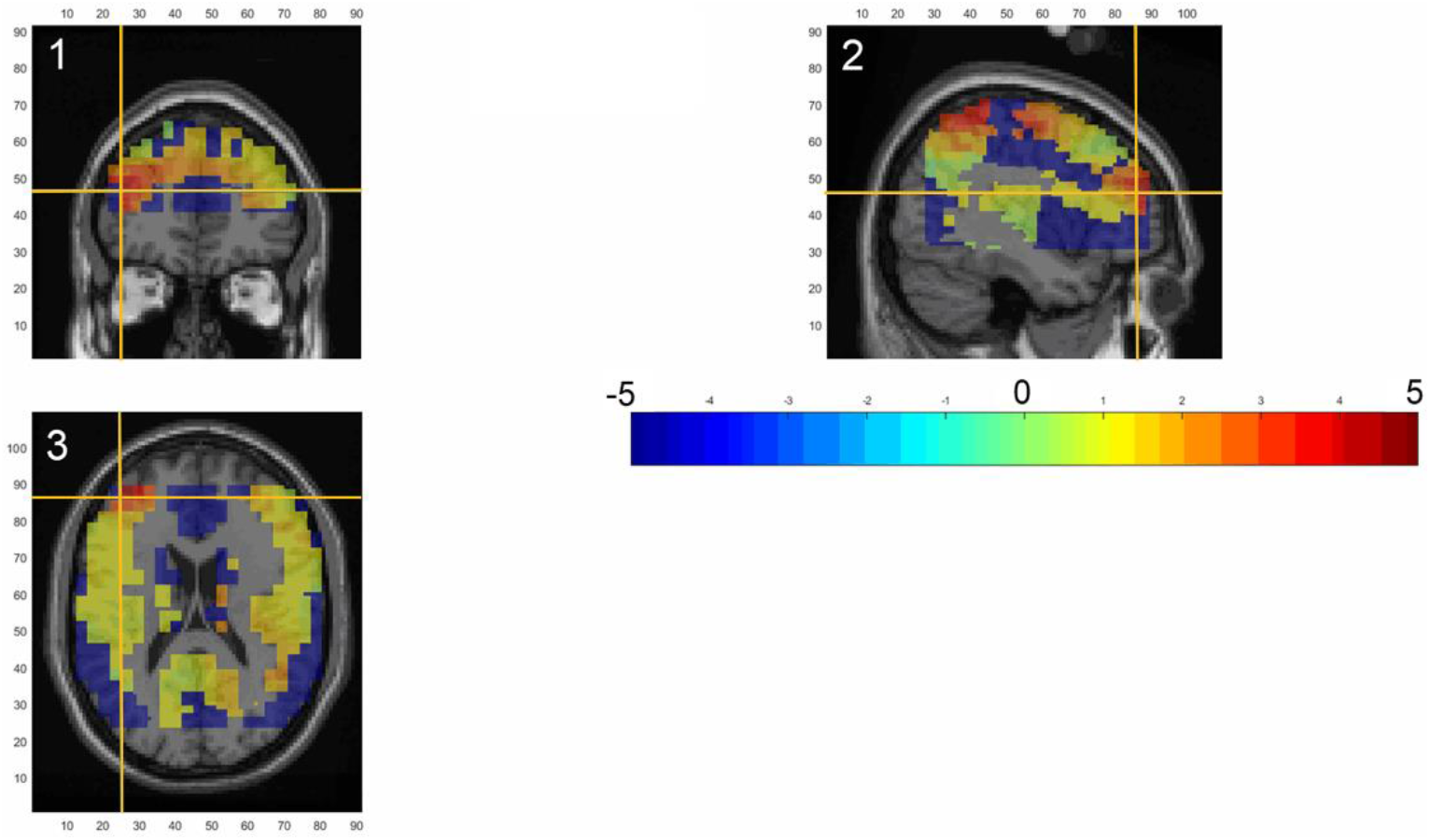
Mean image depicting the source localisation for the inharmonic-harmonic difference ERF activation showing a significant increase in strength for the musician group following 33 Hz entrainment, when an alpha threshold of 0.01 was applied. Location: left middle frontal gyrus (MNI coordinates [− 42 44 18] mm), ~350 ms post stimulus onset, displayed in (1) coronal (2) sagittal and (3) transverse views. Values plotted are the cluster t-statistics within the range −5 to 5 (see colour bar), with significant clusters (*p* < .01) marked by the crosshair.

**Figure 6.**
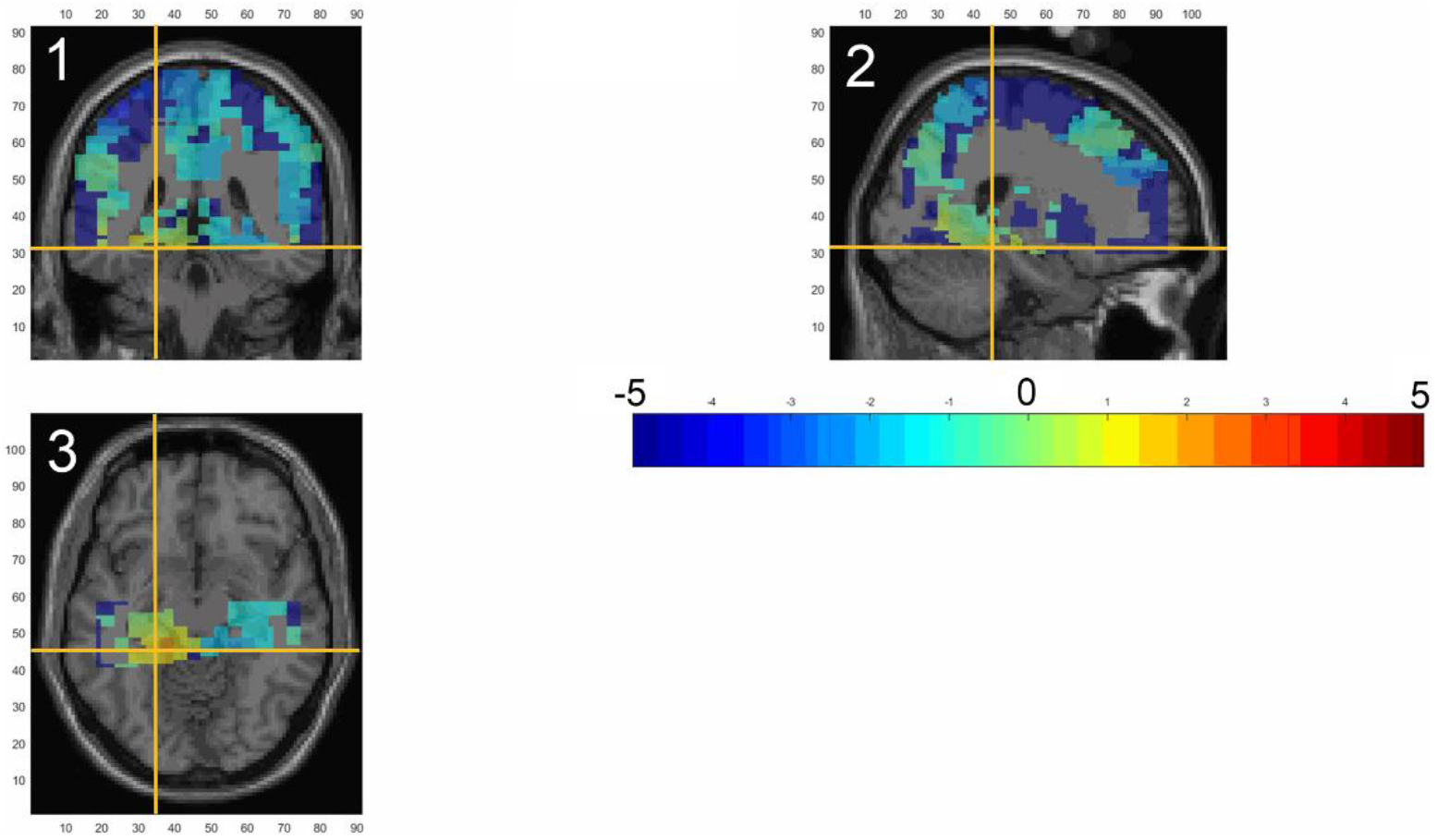
Mean image depicting the source localisation for the inharmonic-harmonic difference ERF activation showing a significant increase in strength for the non-musician group following 33 Hz entrainment. An alpha threshold of 0.01 used. Location: left parahippocampal and left fusiform gyri (MNI coordinates [− 22 −38 −12] mm), ~350 ms post stimulus onset, displayed in (1) coronal (2) sagittal and (3) transverse views. Values plotted are the cluster t-statistics within the range −5 to 5 (see colour bar), with significant clusters (*p* < .01) marked by the crosshair.

Interestingly, in support of the RT findings, which found the responses to the inharmonic were made significantly more quickly than those to the harmonic, for non-musicians, following 33 Hz and 39 Hz entrainment – the results of the source level analysis indicate that a response to a deviant stimulus is stronger for non-musicians compared to musicians, when an aGBR of 33 Hz and 39 Hz is elicited via stimulus entrainment. However, following stimulus entrainment at 33 Hz, RATN source locations differ for musicians and non-musicians. Interestingly, the effects are only evident during the later RATN time window. It is possible that the entrainment stimuli, while containing tonal components, did not elicit an ERAN response which is passively evoked when syntactical anomalies occur in music, or that stimulus entrainment may disrupt the early ERAN response for both groups. However, the suggestion is that the RATN response elicited by irregular tones that are unknown or unpredictable, was improved under certain stimulus entrainment conditions. Considering this convergence of behavioural RT evidence with source activations, a frequency power analysis was conducted to examine the neural evidence for an aGBR evoked through stimulus entrainment using this auditory paradigm, to establish whether the entrainment of an aGBR differed between musicians and non-musicians.

### Frequency power analysis

The stimuli in the experiments conducted by Galambos and colleagues (Galambos et al., 1981; Galambos, 1992) used single pure tones, i.e., comprising only one CF, whereas the current paradigm used two CF’s per pip-train. Furthermore, research using the current paradigm relies on the persistence effects of an evoked aGBR across a range of gamma frequencies. A pure tone burst of 500 ms in duration has been demonstrated to persist within 100 to 120 ms post stimulus onset (Christo Pantev & Elbert, 1994). Furthermore, the current pip-train stimuli have produced persistence effects at up to 400 ms post-stimulus offset (Aksentijevic et al., 2013). Hence, a frequency power analysis was next conducted to explore the neural evidence, at sensor level over temporal cortex, for successful entrainment to the pip-train stimuli that persists during the 100 ms inter-stimulus interval between the prime and the target, dependent on the entrainment frequency, and music ability.

The time frequency data for each entrainment frequency condition, and each group, were epoched for the entrainment period (−1000 ms to 0 ms). A grand average was calculated. These data were used to perform a multi-taper method fast Fourier transform (mtmfft) using a discrete prolate spheroidal sequences (dpss) multi-taper, in Fieldtrip (Oostenveld et al., 2011), to calculate the average frequency content, within the range 28 to 45 Hz, during the 100 ms ISI between the entrainer and target presentation, as measured at all channels covering both temporal lobes, at each entrainment frequency, and for each group. The results were then plotted for each group (see Figure 7).

**Figure 7.**
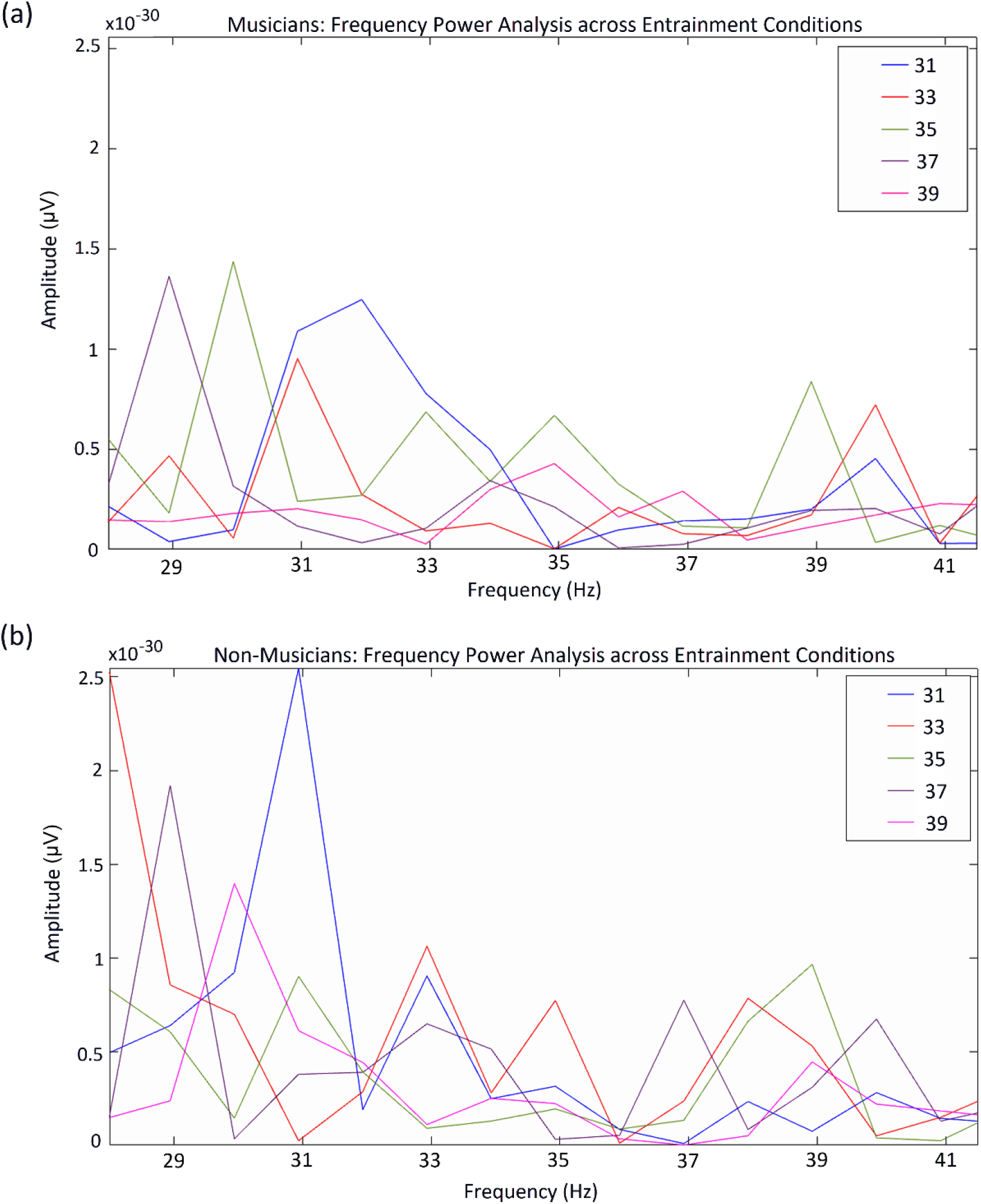
Frequency power spectral analysis during the 100 ms interval between entrainer offset and target onset. (a) Musician; (b) non-musician. Bandpass filtered between 28 and 42 Hz. Legend: Each entrainment condition. Frequency (Hz) is presented on the x-axis and amplitude (μV) is presented on the y-axis.

**Non-musicians:** The amplitude of gamma frequencies between 28 Hz and 31 Hz was much higher for non-musicians. However, and as anticipated, under the 31 Hz, 33 Hz, and 37 Hz entrainment conditions a peak in the corresponding frequency was evident. Furthermore, within the range of 31 Hz to 39 Hz, the highest peak during the 31 Hz, 33 Hz, and 37 Hz entrainment conditions occurs at the corresponding neural oscillatory frequency. This pattern was not evident from the data within the 35 Hz and 39 Hz entrainment conditions, where two peaks occur for frequencies separated by 8-9 Hz. To elucidate, following 35 Hz there was a peak in the amplitude of 31 Hz and 39 Hz, and following 39 Hz entrainment there was a peak in the amplitude of 30 Hz and 39 Hz, and in both cases all other frequencies exhibit low amplitudes. Interestingly, 36 Hz was almost completely absent from all entrainment conditions. **Musicians:** By contrast the power spectrum obtained from the musician group suggested the amplitude of gamma frequencies within the range 31 Hz to 39 Hz was unrelated to entrainment frequency, except perhaps at 31 Hz. However, while a peak in 31 Hz was evident within the corresponding entrainment frequency condition, the trend in the data for this condition, and similarly for an entrainment frequency of 33 Hz suggested a more exaggerated two-peak pattern, with a spike in the amplitude of 31-32 Hz and 40 Hz, and a complete attenuation of 35 Hz, hence a peak-to-peak separation of 9 Hz.

Thus, the evoked aGBRs demonstrated signs of a 9 Hz (alpha) modulation, which was more pronounced for musicians. Furthermore, for non-musicians, following each entrainment the greatest peak in amplitude occurred for the corresponding frequency, suggesting the aGBR was successfully evoked. Given the 8-9 Hz separation between the frequencies that exhibit higher amplitudes, under some entrainment conditions, in the 100 ms period between the termination of the entrainment stimulus and the onset of the target, a phase-amplitude coupling analysis was conducted to explore underlying interactions which might give rise to this effect.

### Phase-amplitude coupling (PAC) results

In addition to the effect of priming on RT responses to a deviant stimulus, found for non-musicians, findings from previous work strongly suggests that enhanced (i.e., significantly faster) RT responses to deviant tones are a consequence of an interaction between the aGBRs, evoked by the entrainment stimulus, and endogenous brain rhythms in the theta range – approximately 2 Hz below the suggested influence of alpha on the evoked aGBRs, inferred from the spectrograms in Figure 7 (Elliott, 2014; Elliott & du Bois, 2017). The current research provided the opportunity to investigate the support for such an interaction through an examination of gamma-theta phase-amplitude coupling (PAC), whereby the instantaneous phase of endogenous theta modulates the instantaneous amplitude of gamma oscillations. PAC coupling has been linked to working memory processing and information exchange by providing windows of coherence between participating neural networks (Penny, Duzel, Miller, & Ojemann, 2008; Scheffer-Teixeira et al., 2012; Schomburg et al., 2014). Figure 8 to Figure 10, and Supporting Figure 1 to Supporting Figure 5 in supporting material, illustrate the modulation of the amplitude of gamma frequencies covering the range of entrainment frequencies used in this paradigm by the phase of high delta, theta, and alpha during each entrainment condition, for both groups. The averaged PAC values for each group, are presented in the top row of these figures, marked (a). The darker the colour of the voxels in these colour comodulograms, the higher the PAC value for that frequency combination, as can be seen from the indictor at the right side of the comodulograms. When interpreting the gray scale comodulograms in the second row, marked (b), the grey and white voxels represent the proportion, or percentage, of the sample for whom this frequency combination had a PAC value that was significant at alpha levels of 0.05 and 0.01, respectively, as can be seen by the indictor at the right side of the comodulograms, i.e., black voxels were not significant and the number at the top of the column (indicator) represents the proportion of people for whom the PAC was significant. By comparing the peaks in the spectral density graphs, marked (c) for the slow waves, and (d) for the gamma frequencies, with these comodulograms ((a) and (b)), consistencies between high PAC values that were also found to be significant, and the proportion of that group (musician or non-musician) for whom they were significant, can be examined.

**Figure 8.**
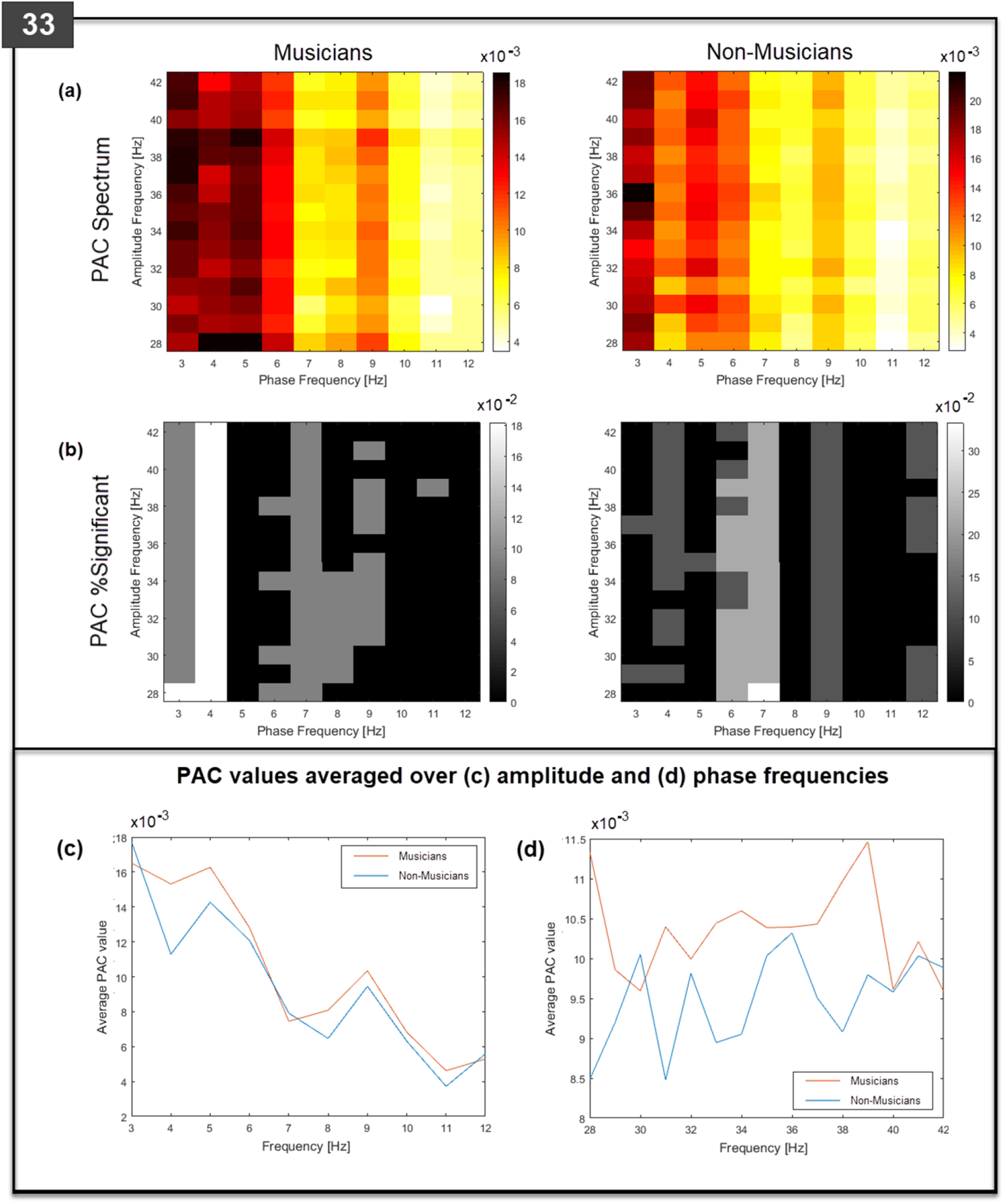
Grand-averaged PAC spectral densities for musicians and non-musicians during the 33 Hz entrainment condition (top left corner). (a) Grand-averaged PAC values for each gamma and slow wave frequency combination in the spectrum. (b) Percentage cases with significant PAC (white; *p* < 0.01, and grey; *p* < 0.05) for each gamma and slow wave frequency combination in the spectrum. (c) and (d) Spectral densities for ‘musicians’ (red legend) and ‘non-musicians’ (blue legend).

**Figure 9.**
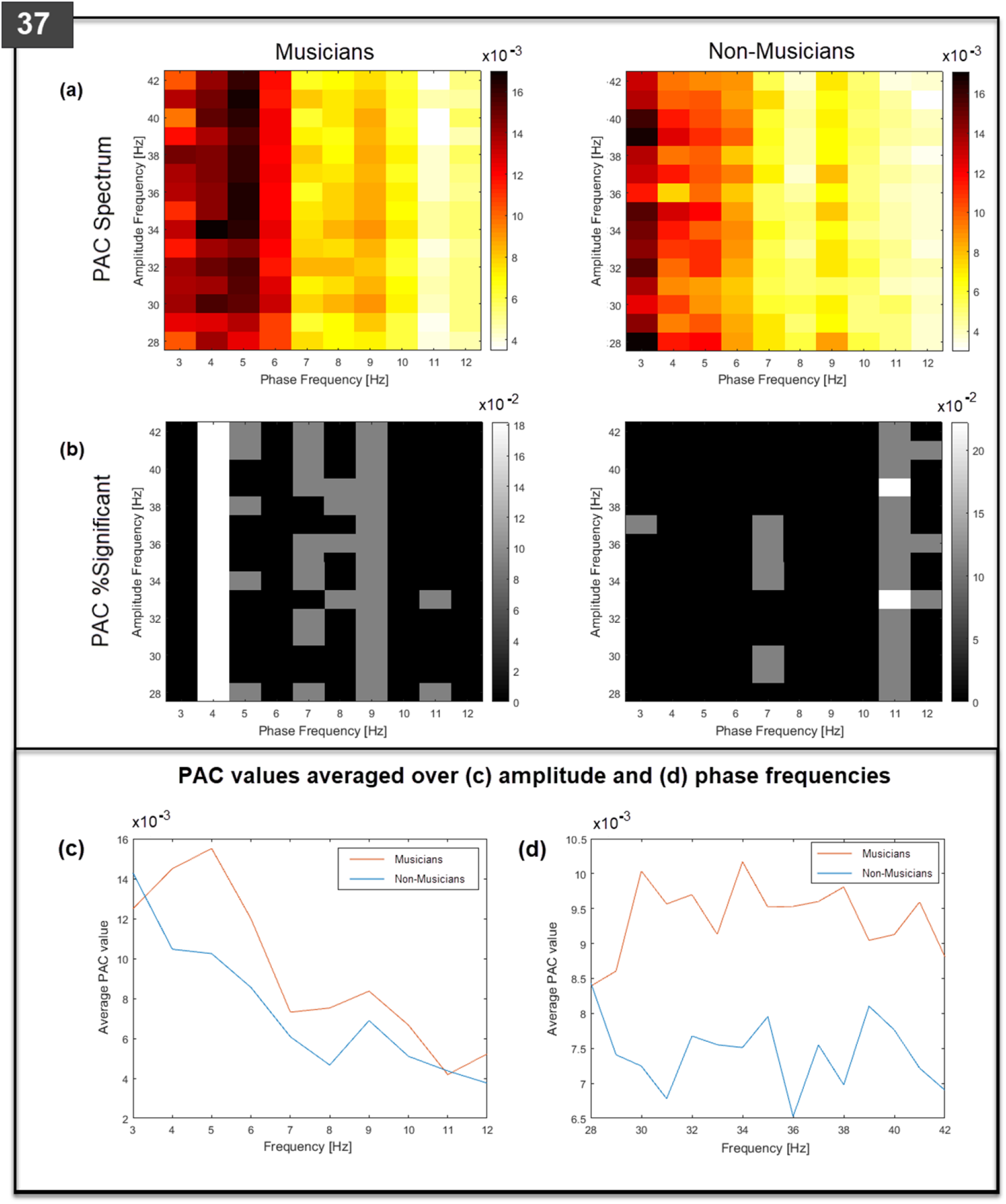
Grand-averaged PAC spectral densities for musicians and non-musicians during the 37 Hz entrainment condition (top left corner). (a) Grand-averaged PAC values for each gamma and slow wave frequency combination in the spectrum. (b) Percentage cases with significant PAC (white; *p* < 0.01, and grey; *p* < 0.05) for each gamma and slow wave frequency combination in the spectrum. (c) and (d) Spectral densities for ‘musicians’ (red legend) and ‘non-musicians’ (blue legend).

**Figure 10.**
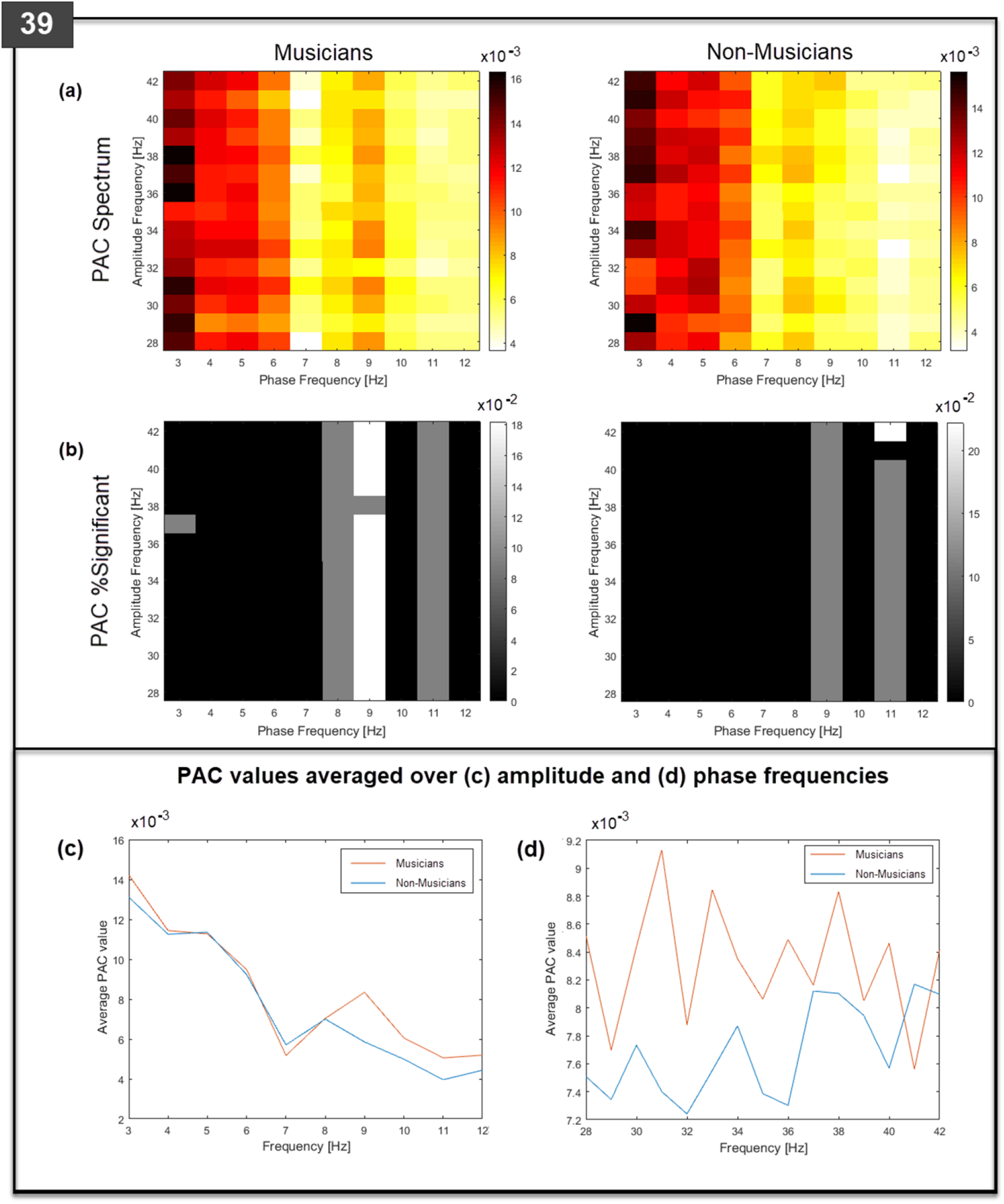
Grand-averaged PAC spectral densities for musicians and non-musicians during the 39 Hz entrainment condition (top left corner). (a) Grand-averaged PAC values for each gamma and slow wave frequency combination in the spectrum. (b) Percentage cases with significant PAC (white; *p* < 0.01, and grey; *p* < 0.05) for each gamma and slow wave frequency combination in the spectrum. (c) and (d) Spectral densities for ‘musicians’ (red legend) and ‘non-musicians’ (blue legend).

A within-subject analysis of changes in PAC across entrainment conditions found a significant difference for non-musicians only, between the 33 Hz and 37 Hz entrainment conditions (see Figure 8 and Figure 9). It is speculated that this is driven by a 6 – 7 Hz theta modulation that is significant for 25 – 30 % of the non-musician group following entrainment of a 33 Hz GBR compared to entrainment of a 37 Hz GBR. Interestingly, 36 Hz appears to be attenuated, i.e., not present, across entrainment conditions in the spectrogram for non-musicians (see Figure 7).

Notably, the proportion of musicians contributing to a significant PAC value does not exceed 18% of the overall group – yet remains consistent across conditions and seems to be generally representative of high pac (r_pac_) values (see the indicator on the right of the significance comodulograms for musicians, in Supporting Figure 1 to Supporting Figure 5 in supporting material). Moreover, the peaks in the spectral density graphs are largely representative of the phase modulation frequencies that produce a modulation effect, i.e., have higher PAC values, across the spectrum of amplitude frequencies, for approximately 18% of the sample, in each condition. The most prominent frequency modulations occur in phase with 4 Hz theta, and 8-9 Hz alpha rhythms. Interestingly, there is a complete absence of a 4 Hz modulation effect under the 39 Hz entrainment condition, coupled with lower averaged delta range PAC values across the gamma amplitude frequencies, evidenced for both groups.

In contrast, more variation occurs between conditions for the proportion of non-musicians for whom the same frequency combinations resulted in significant PAC values – as can be seen from the indicator to the right of the significance comodulograms, for each entrainment condition, for non-musicians; 31 Hz (10%), 33 Hz (30%), 35 Hz (20%), 37 Hz (20%), and 39 Hz (20%) (see Supporting Figure 1 to Supporting Figure 5 in supporting material). Thus, the 4, 9 and 12 Hz modulation during the 31 Hz entrainment condition, which was observed for all amplitude frequencies, was only significant for 10% of the group, while the 7 Hz modulation during the 33 Hz entrainment condition was significant across the full range of gamma for 30%+ of the non-musicians, and a 9 Hz modulation across the full gamma range for ~15-20%. While not extending across the full range of gamma, the 6 Hz modulation was also representative of 30%+ of the group – and corresponding PAC values were slightly increased in strength compared to the 7 and 9 Hz interactions, as can be seen from the spectral density graph (c) in Figure 8. Interestingly, previous research using stimulus entrainment in both the visual and auditory domain has asserted that a facilitation of responses, notably the pop-out effect associated with 33 Hz entrainment, can be attributed to an interaction with a theta rhythm of approximately 6-7 Hz (Aksentijevic et al., 2011; Elliott, 2014; Elliott & du Bois, 2017). The significant 6-7 Hz modulation observed in these data for a approximately one-third of the sample following 33 Hz entrainment, supports this assertion – and may reflect a greater influence of entrainment on non-musicians’ tone discrimination. In turn, this observed influence suggests a greater reliance on the stimulus (bottom-up) information, that is subserved by gamma-band frequencies, while the more general phase modulation of gamma amplitude by specific endogenous frequencies found for musicians may echo a top-down training induced response.

## Discussion

The current research aimed to investigate functional differences in the neural mechanisms of auditory cognition that derive from music training. The purpose of using an auditory prime that simultaneously primes a harmonic expectancy (based on the carrier frequencies) and entrains a gamma-band response that is phase locked to the primed AM frequency, was to provide a means to examine these functional differences through the lens of the priming effect – and how or if this effect differs when the brain has been plastically altered through music training.

Previous research has shown differences in priming the auditory system with gamma-band entrainment frequencies that occur as a consequence of music training (Aksentijevic et al., 2011, 2013, 2014). An aGBR is presumed to be evoked by, and phase-locked to, the entrainer stimulus. External stimulus information is known to be carried by frequencies in the gamma-band range as the phase cycle of these frequencies is fast enough to feedforward prediction errors to update models that have been based on previous learning (Fries, 2015; Michalareas et al., 2016). Moreover, the harmonic relations embedded in the entrainer stimulus generates the listeners’ expectancies regarding the next event – the harmonic target is an expected event (consonant), and the inharmonic is an unexpected, or deviant event (dissonant). Consistent with previous research, the results from the behavioural (RT) data revealed a 33 Hz entrainment frequency specific pop-out effect for non-musicians only. However, deviating from previous findings – a pop-out effect was also found following 39 Hz entrainment (see Figure 2). Furthermore, the musicians in the current sample did not exhibit faster RTs compared to non-musicians – possibly a reflection of differences in sensitivity to auditory outliers due to type of music training (current classical versus previous jazz, musicians). Importantly, consistent with the Aksentijevic sample, musicians’ responses to inharmonic targets, while faster than responses to harmonic targets, did not differ significantly across all rates of entrainment (inharmonic-harmonic RTs, *p* > 0.05) (Aksentijevic et al., 2014).

In general, differences in the effect of priming using stimulus entrainment as a function of music training were demonstrated in relation to the RATN response – an evoked response to irregular tone structures within an unpredictable sequence (Headley & Weinberger, 2011). Source analyses during this later (RATN) time window (325-375 ms), found the ERF response to the inharmonic (deviant stimulus) to be significant across both groups following 33, 37, and 39 Hz entrainment, consistent with the finding that the RT differences between the harmonic and inharmonic targets reached significance for these entainment conditions (see Figure 3). Sources were located in the left hippocampus and parahippocampus (33 Hz), right superior frontal gyrus (SFG) (37 Hz), and the left inferotemporal gyrus (ITG) (39 Hz). In line with significant RT differences, the source of these harmonic ERF differences following 33, and 39 Hz entrainment only remained significant for non-musicians, when the data were analysed separately per group (see Figure 4).

The right SFG has been associated with efficient response inhibition, characterising the neural processes associated with estimating likelihood – referred to as conflict anticipation (Hu, Ide, Zhang, & Li, 2016). Furthermore, activation of this brain region has been implicated in improved auditory working memory performance (Othman et al., 2019). Thus, activation of the right SFG following 37 Hz entrainment suggests that the effect of priming at this rate facilitates response inhibition, and overall performance, for both groups. While grey matter density of the hippocampus has been found to be higher for musicians and musical expertise induces hippocampal structural and functional plasticity (M Groussard et al., 2009; Mathilde Groussard et al., 2010; Herdener et al., 2010), the hippocampus has also been found to be involved in novelty detection (Kumaran & Maguire, 2007a, 2007b). Operating as a comparator, the hippocampus generates a response when the incoming stimulus violates predictions based on previous learning. Moreover, hippocampal activity is related to detection of novel configurations of familiar items – referred to as associative novelty (Kumaran & Maguire, 2007a). Furthermore, activation of the left ITG among non-musicians following 39 Hz entrainment is possibly related to the contrast between the inharmonic and harmonic auditory stimuli given this area of cortex similarly exhibits increased activation in response to pseudowords compared to words (Binder, 2000; Specht & Reul, 2003). The rationale for a contrast related response is based on the consideration that non-musicians process the pip-trains based on the contrast between the tones, rather than the harmony relations.

Critically, a source level ERF difference following 33 Hz stimulus entrainment uncovered differences in source activation strengths for each group – in separate locations, thus, highlighting a differential effect of priming on functional connectivity that is dependent on music training. For non-musicians the source remained located in the left hippocampus and parahippocampus regions – consistent with the enhanced ERF difference between source activations in response to the inharmonic compared to the harmonic found for this group alone, following 33 Hz stimulus entrainment (see Figure 5). For musicians the location of the increased difference ERF was in the left middle frontal gyrus (MFG) (see Figure 6). As discussed, hippocampal activation relates to novelty detection. Specifically, left hemispheric hippocampal activity occurs when one component of a familiar sequence is novel, while another is repeated (Kumaran & Maguire, 2007a). In the context of the current paradigm, the implication is that for the musically naïve brain, 33 Hz entrainment facilitates a response to the inharmonic (a novel stimulus in an otherwise repeated sequence of pips). Musicians on the other hand, as a consequence of their training, have developed more efficient neural strategies, which may explain the increased activation originating in the left MFG for musicians compared to non-musicians, observed following 33 Hz entrainment. The right MFG is associated with working memory processes and storage in general, with musicians showing increased activation compared to non-musicians under increased load (Leung, Gore, & Goldman-Rakic, 2002; Olesen, Westerberg, & Klingberg, 2004; Pallesen et al., 2010). However, musicians have demonstrated enhanced functional connectivity across a neural network that spans between the left primary auditory cortex and the right globus palidus, bilateral motor areas and supramarginal gyri (SMG), left premotor cortex, and the left MFG (Fauvel et al., 2014). Thus, it is suggested that the 33 Hz entrainment improves musicians’ recruitment of learning induced working memory neural strategies.

Our source level analysis suggests functional differences in auditory cognition between musicians and non-musicians are separable through an examination of differences in the effect of priming. Notably, stimulus entrainment appears to directly facilitate non-musicians’ responses to the primed stimuli, whereas the only benefit of entrainment for musicians is suggested to be through an enhancement of an existing, training induced, functional connectivity network. Further support for this interpretation was provided by the frequency power analysis and the phase-amplitude coupling analysis. Stimulus entrainment entails driving brain oscillations using an external periodic force and is characterised by the adjustment of phase to the external stimulus (phase-locking) and an increase in amplitude among participating neurons (Thut, 2011). Therefore, an amplification of entrainment frequencies in auditory cortex was examined by plotting the spectrogram, covering the range of entrainment frequencies, during the 100 ms period between entrainment and target presentation. Consistent with stimulus entrainment, following 31, 33, and 37 Hz entrainment, the anticipated peak in the corresponding frequency was evident in the spectrogram for non-musicians. However, the spectrograms of musicians presented a much more complicated pattern of frequencies for each entrainment condition that imply an effect of a 9 Hz modulation during the 31 and 33 Hz entrainment conditions (see Figure 7). Furthermore, a phase-amplitude 8-9 Hz modulation effect was found consistently for musicians, in terms of the percentage of the group for whom the effect was significant, as well as across all conditions. The range and consistency of the phase-amplitude coupling evident for the non-musicians fluctuates in contrast.

Notably, the 7 Hz phase modulation of gamma amplitude across the full range, was found to be significant for between 27 and 33% of non-musicians following 33 Hz entrainment (see Figure 8). The first research using the current auditory priming paradigm, and previous research using a functionally similar visual priming paradigm which used rates of flicker to evoke a phase locked cortical frequency response, have consistently supported the suggestion that an effect of priming, i.e., faster reaction-time responses following specific entrainment rates, depended on a phase interaction with a slower endogenous theta rhythm, of 6.69 Hz (see Elliott & Müller, 1998 for details of the visual priming paradigm) (Aksentijevic et al., 2011; Elliott, 2014; Elliott & du Bois, 2017; Elliott & Müller, 2004; Elliott & Müller, 1998). The results reported here represent the first evidence at the cortical level in support of this inference. The accumulated evidence from the source analyses, the frequency power spectrograms, and the PAC analyses infer a reduced effect of stimulus entrainment for musicians that stems from a greater reliance on the modulation effects of spontaneous theta and alpha on gamma-band synchronisation, rather than the early entrainment of gamma-band information carriers. In other words, musicians have developed top-down neural strategies that supersede the effects of stimulus entrainment – consistent with previous support for the redundancy of gamma-band oscillatory influence on musicians’ performance due to experience-dependent plasticity (Headley & Weinberger, 2011).

The selection of relevant sensory input requires theta modulated synchronised gamma-networks, whereby cycles of spontaneous theta make and break the synchronised gamma oscillations through a process of phase reset (Fries, 2009; Rollenhagen & Olson, 2005). Research has revealed that neurons in the IT cortex of monkeys respond selectively to a preferred visual stimulus from a display of multiple stimuli through this alternating pattern of rhythmic theta modulation, characterised by an increase in firing rate in response to a preferred stimulus and a decrease in firing rate in response to a non-preferred stimulus (Rollenhagen & Olson, 2005). Moreover, the firing rate was enhanced when the preferred stimulus was presented in the presence of the non-preferred stimulus - interesting given the 2000 and 2400 Hz tones of the harmonic and inharmonic targets in the current research were presented simultaneously with the baseline 1000 Hz tones. In the context of this research a further relevant consideration has been suggested by Fries (2009), which is that this rhythmic response to the preferred stimulus was consistent with a 5 Hz theta oscillation, allowing for the selection of a response every 200 ms (1000/5 = 200), and therefore when two stimuli need to be discriminated, a selection is required every 100 ms, corresponding with a 10 Hz alpha rhythm, also found to modulate gamma-band synchronisation (Fries, 2009). While the IT cortex is critical for visual object recognition, as previously noted activity in IT has also been associated with contrast recognition in the auditory domain (Binder, 2000; Specht & Reul, 2003).

Although the thalamus is fundamentally involved in the generation of alpha (Roux, Wibra, Singer, Aru, & Uhlhaas, 2013), with theta generation occurring in the hippocampus and hippocampal structures, (Pignatelli, Beyeler, & Leinekugel, 2012), alpha has also been recorded in the hippocampus and theta in the frontal cortex (Buffalo, Fries, Landman, Buschman, & Desimone, 2011; Roux & Uhlhaas, 2014) – and both have been associated with working memory (WM). According to the short-term memory model proposed by Horn and Usher (1992), stimulus information is carried forward within a synchronised gamma-band oscillatory network, with sequential stimuli coded by sequential gamma subcycles. These subcycles are nested in each theta cycle, allowing sequence order to be coded and stored in short-term memory, which is supported by the well-established 7 ± 2 short-term memory capacity limit (Horn & Usher, 1992; Jensen & Lisman, 2005). Furthermore, whereas theta-gamma modulation is involved in WM item maintenance, alpha-gamma modulation has been found to protect items in WM through the inhibition of non-relevant information (Roux & Uhlhaas, 2014). Interestingly, theta phase modulation of alpha amplitude has been found to be involved in attending to multiple items simultaneously (Song, Meng, Lin, Zhou, & Luo, 2014). Furthermore, enhanced theta activity has been demonstrated to be associated with prediction, i.e. when a prime accurately predicts a response (Dikker & Pylkkänen, 2013). These frequency interactions, taken together with the enhanced ERF difference for inharmonic targets following a 37 Hz entrainment frequency, highlight an important effect of priming that spans both groups – that is, rather than facilitating auditory cognitive processes alone, priming at some entrainment frequencies yields an enhancement of cognitive processes elicited by task demands, i.e., the requirement to make a forced-choice response, as quickly as possible, based on the tonal difference between two targets. However, it is important to note, that real-world auditory cognition does require selective and sustained attention, which in turn requires inhibition of irrelevant information – and importantly, to navigate effectively and interact with the environment, auditory cognition (and cognition in general) requires the ability to correctly predict events as they unfold. This latter requirement is fundamental to a musician’s ability to synchronise their performance with other musicians.

It is also worth noting, within the auditory cortex nested oscillations are hierarchically organised such that the phase of 1–4-Hz neural oscillators modulate the amplitude of 4–10-Hz neural oscillators, which in turn modulate gamma amplitude (30-50 Hz), thus optimising the system’s ability to process rhythm through structuring the pattern of temporal activity (Lakatos et al., 2005). Thus, in tandem with the benefits associated with the modulation effects of spontaneous brain rhythms on cognition in general, auditory cognition relies on this hierarchy of interactions to track and separate sound objects. Furthermore, it has been argued that sensory and cognitive events evoke superimposed delta, theta, alpha, and gamma oscillatory mechanisms, within selectively distributed networks, varying in intensity, synchronisation, delay and duration, depending on functional demands (Başar, Başar-Eroglu, Karakaş, & Schürmann, 2001).

## Conclusion

In summary, while both groups exhibit behavioural (RT) and ERF activity related to priming an auditory response using stimulus entrainment presented at entrainment frequencies of 33 Hz, 37 Hz, and 39 Hz – there was a disparity in the outcome of this effect dependent on music ability which was found to reflect functional differences in auditory cognition as a consequence of music training. Notably, non-musicians demonstrated a direct enhancement in working memory processes and ability to detect a deviant auditory stimulus, whereas the effect for musicians reflected improved functional connectivity, emphasised by consistent theta and alpha phase modulation of gamma amplitude, presumably due to the plasticity effects of music training. These slower endogenous frequencies are relied upon for feedback predictions, e.g. inhibiting irrelevant information (via alpha rhythms) and manipulating information (via theta rhythms) (Kirmizi-Alsan et al., 2006; Roux & Uhlhaas, 2014; Sauseng, Griesmayr, Freunberger, & Klimesch, 2010; Sauseng et al., 2009). Thus, it is concluded from the findings of this research that musicians’ response to the harmonic structure of a sound depends more on top-down processing (greater range of interactions with slower brain rhythms) than bottom-up processing (gamma rhythms) provided by the entrained prime, found to facilitate the responses of non-musicians.

## Limitations and future research

The current analysis provides motivating information regarding the utility of stimulus entrainment in an examination of differences in auditory cognitive processes dependent on musical ability. However, further analysis is recommended using a larger sample to improve the power of the statistical findings. Recent literature examining structural and functional differences between musicians and non-musicians using MEG have generally had a sample size of 15, and up to 26, per group. A power analysis for an effect size of 0.25, at 80% power, advises 17 per group. Thus, the current study was underpowered. Furthermore, towards a more complete understanding of the importance of gamma frequency-specific effects on auditory cognition, a wider range of entrainment frequencies should be employed. A further consideration that has been highlighted here is the effect of music training type. It would be interesting to employ the same design using a sample of jazz musicians to compare the results. Regarding the range of expertise within the current sample, varying from grade 3 to professional – this represents a further limitation which should be addressed in future work.

## acknowledgements

This work was supported by an Irish Research Council scholarship to the first author (project ID: GOIPG/2015/1678), and in part by the Northern Ireland Functional Brain Mapping (NIFBM) Facility Project through Invest Northern Ireland (Invest NI) and the Ulster University under Grant 1303/101154803. We thank Dr Pramod Gaur (Ulster University, Magee campus, Derry) for assisting in data collection, Dr Tamas Minarik and Prof Paul Sauseng (Ludwig-Maximilians-Universität Munich) for contributing to the analysis codes, and Dr Bernadette C. M. van Wijk for advising on the phase-amplitude coupling analysis (The Wellcome Trust Centre for Neuroimaging, UCL, London).

## Data availability statement

The data that support the findings of this study are available from the corresponding author upon reasonable request.

