## Supporting Material for "Functional Differences in the Neural Substrates of Auditory Cognition as a Consequence of Music Training"

### Annex 1: Supporting materials

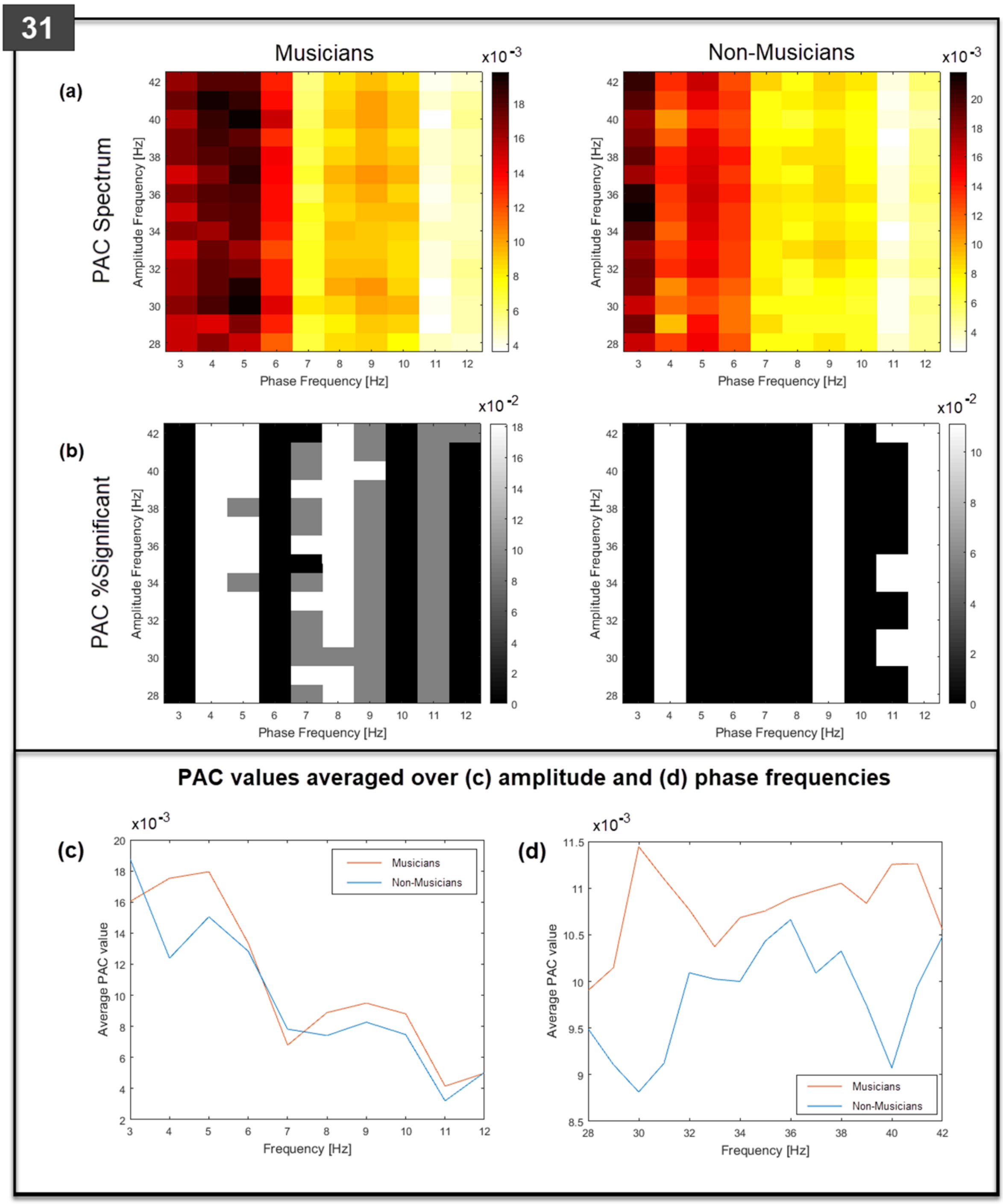


**Supporting Figure 1. Grand-averaged PAC spectral densities for musicians and non-musicians during the 31 Hz entrainment condition (top left corner).** (a) Grand-averaged PAC values for each gamma and slow wave frequency combination in the spectrum. (b) Percentage cases with significant PAC (white; $p < 0.01$, and grey; $p < 0.05$) for each gamma and slow wave frequency combination in the spectrum. (c) and (d) Spectral densities for 'musicians' (red legend) and 'non-musicians' (blue legend).

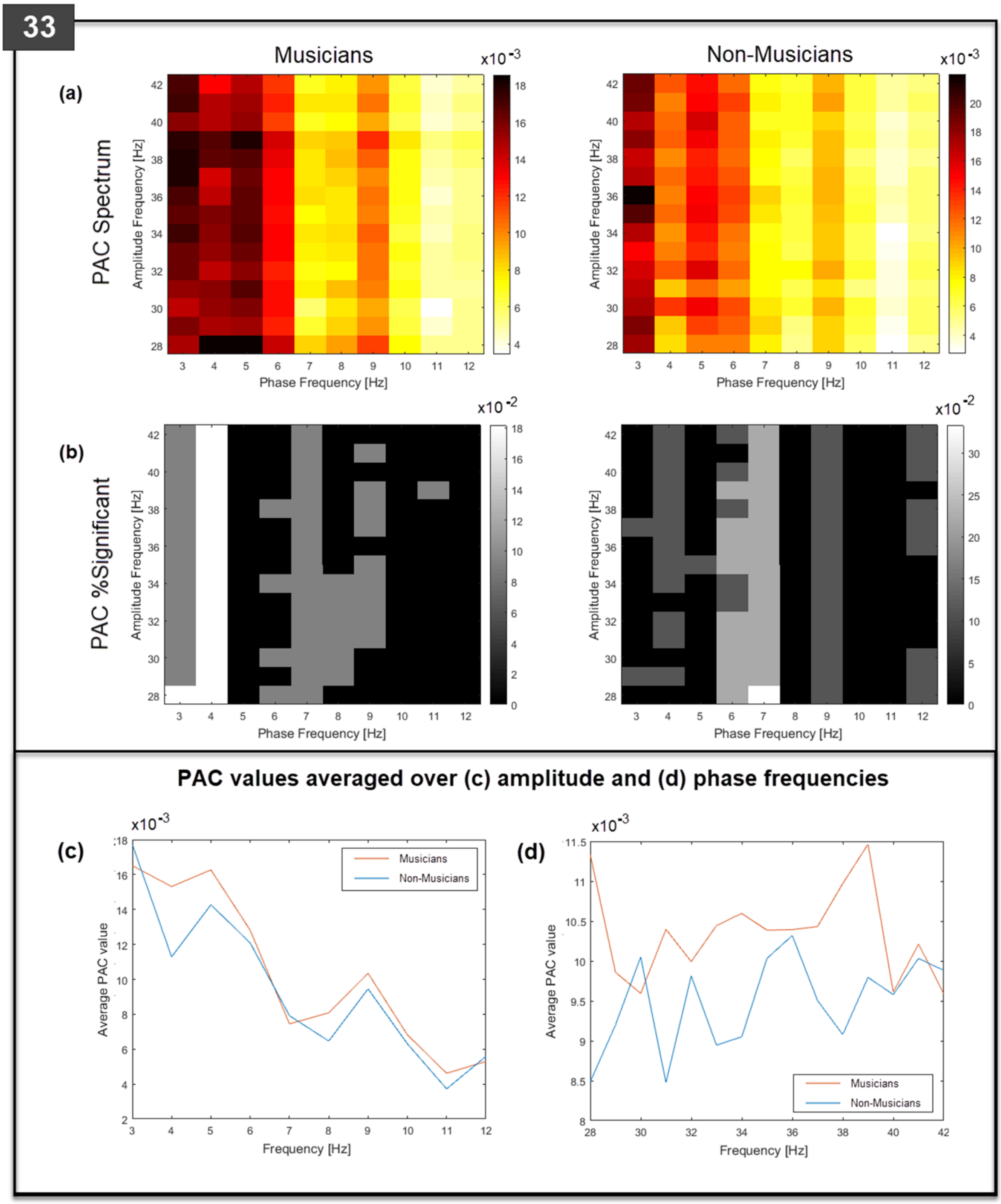


**Supporting Figure 2. Grand-averaged PAC spectral densities for musicians and non-musicians during the 33 Hz entrainment condition (top left corner).** (a) Grand-averaged PAC values for each gamma and slow wave frequency combination in the spectrum. (b) Percentage cases with significant PAC (white; $p < 0.01$, and grey; $p < 0.05$) for each gamma and slow wave frequency combination in the spectrum. (c) and (d) Spectral densities for 'musicians' (red legend) and 'non-musicians' (blue legend).

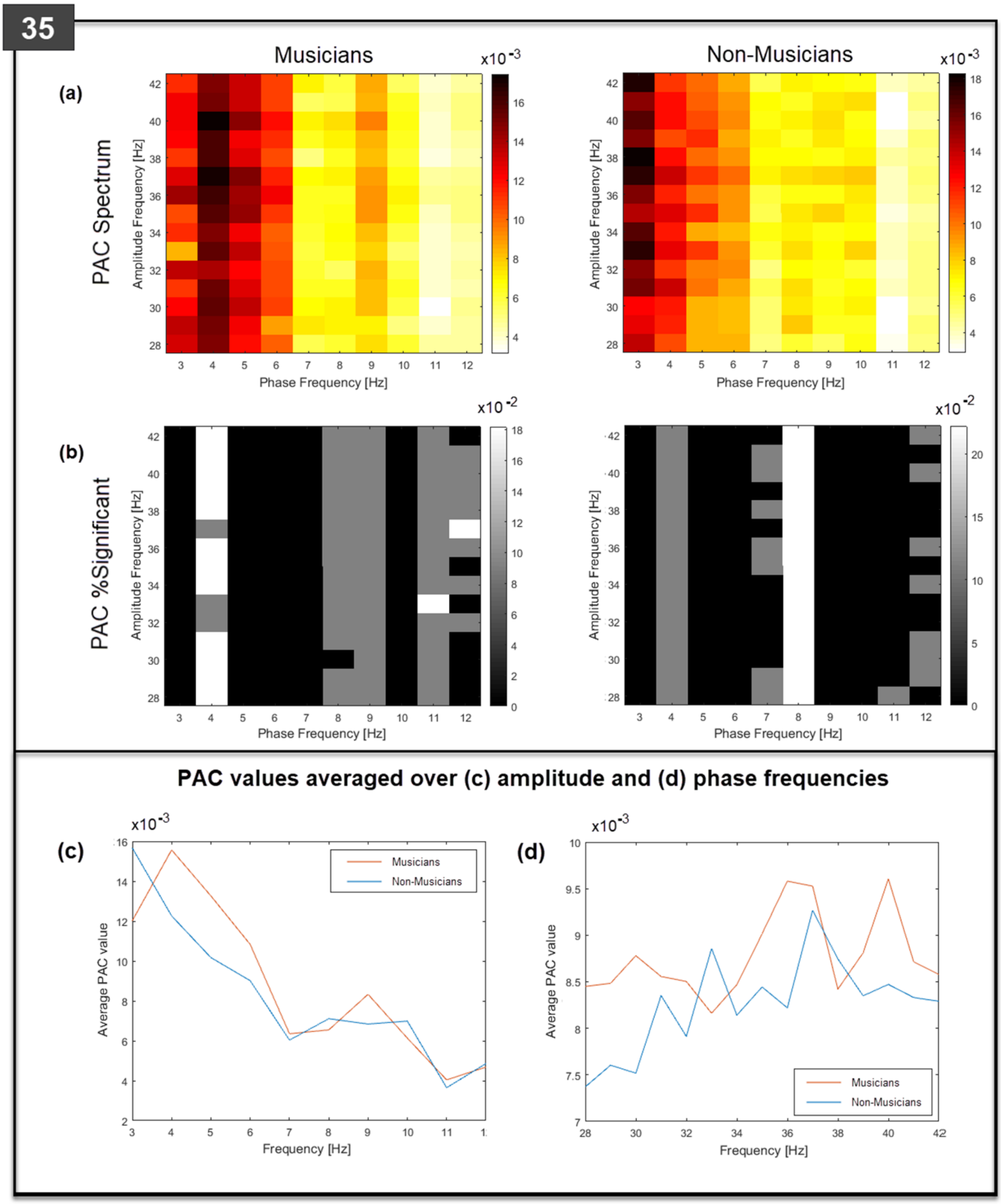


**Supporting Figure 3. Grand-averaged PAC spectral densities for musicians and non-musicians during the 35 Hz entrainment condition (top left corner).** (a) Grand-averaged PAC values for each gamma and slow wave frequency combination in the spectrum. (b) Percentage cases with significant PAC (white; $p < 0.01$, and grey; $p < 0.05$) for each gamma and slow wave frequency combination in the spectrum. (c) and (d) Spectral densities for 'musicians' (red legend) and 'non-musicians' (blue legend).

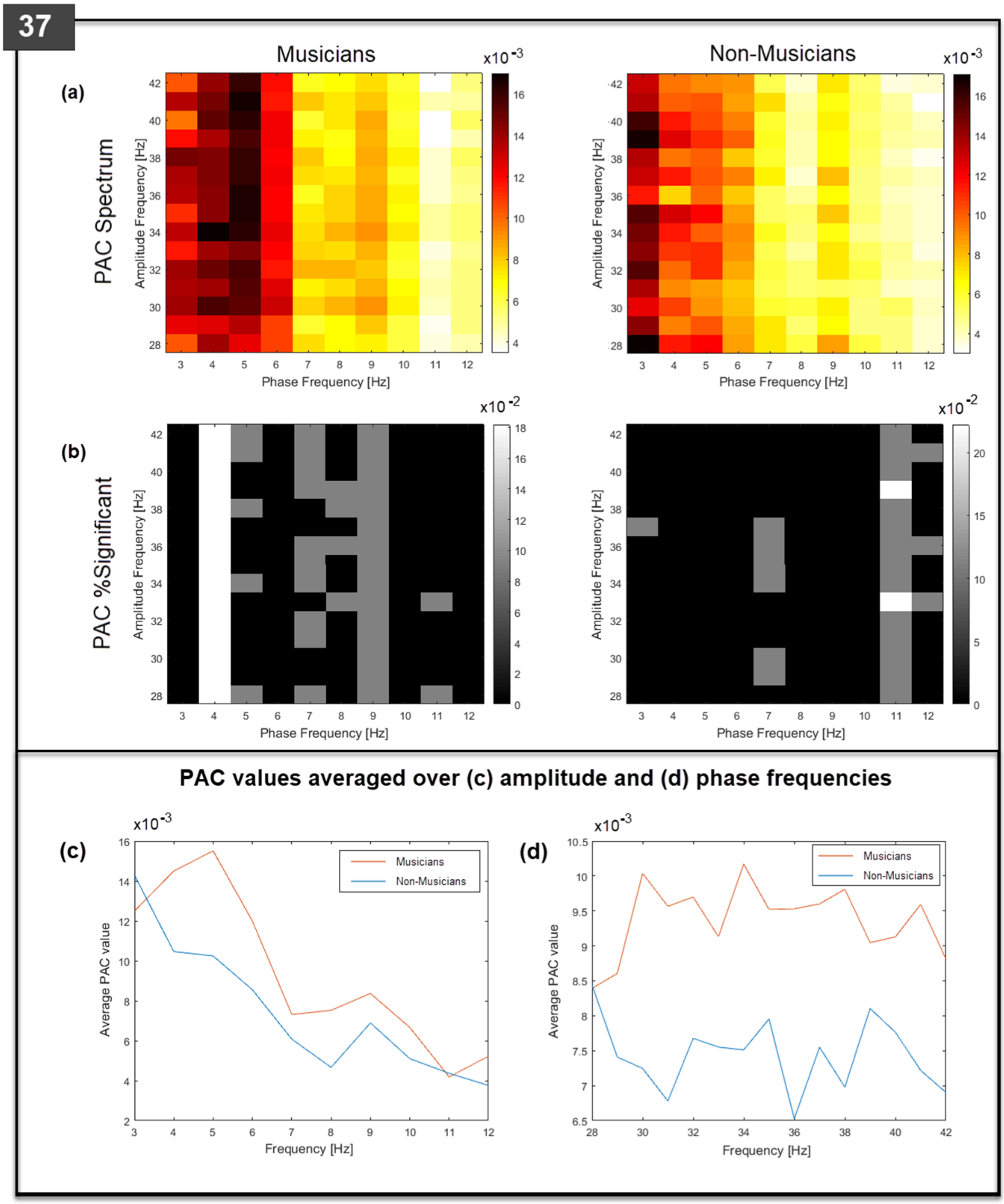


**Supporting Figure 4. Grand-averaged PAC spectral densities for musicians and non-musicians during the 37 Hz entrainment condition (top left corner).** (a) Grand-averaged PAC values for each gamma and slow wave frequency combination in the spectrum. (b) Percentage cases with significant PAC (white; $p < 0.01$, and grey; $p < 0.05$) for each gamma and slow wave frequency combination in the spectrum. (c) and (d) Spectral densities for 'musicians' (red legend) and 'non-musicians' (blue legend).

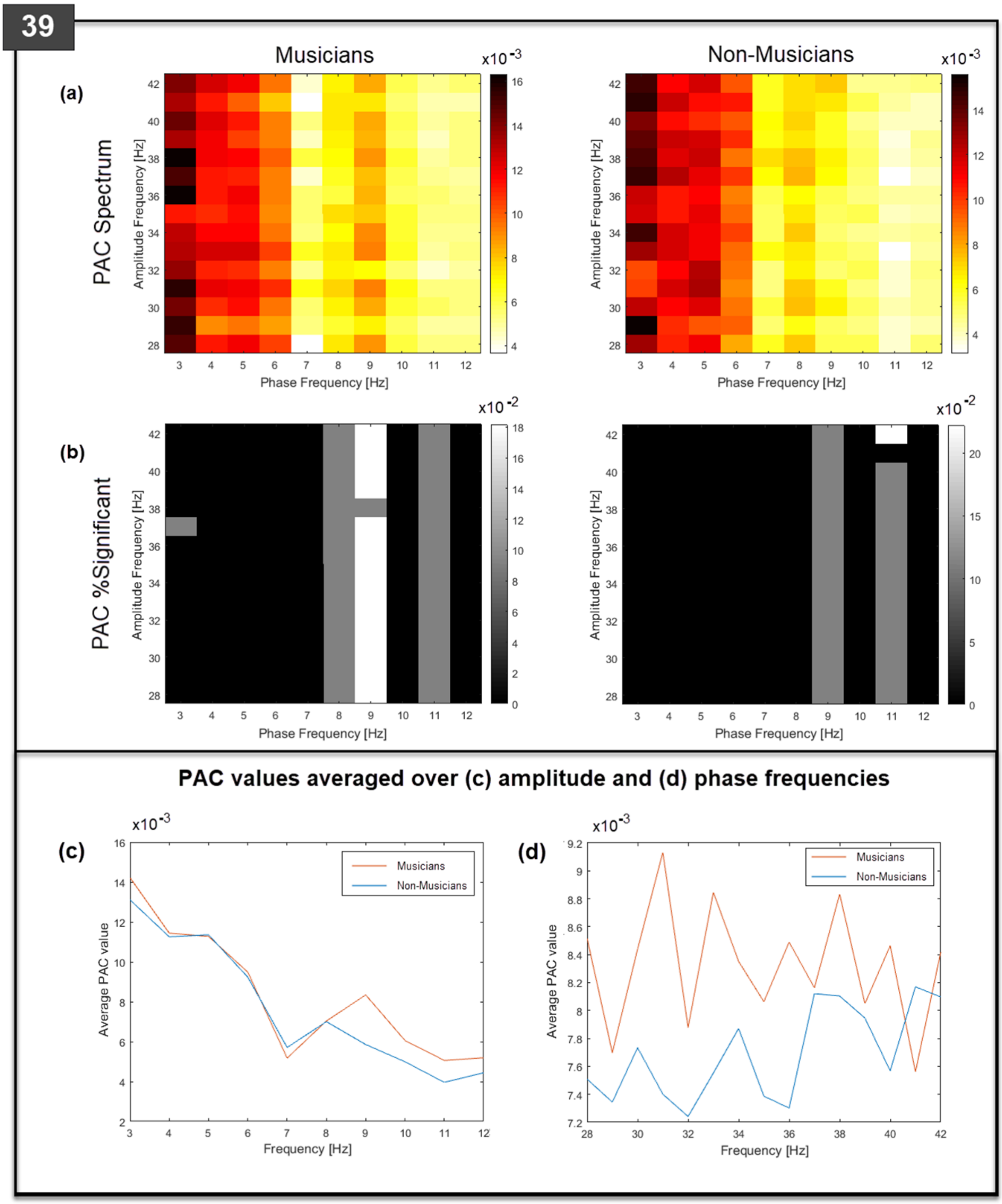


**Supporting Figure 5. Grand-averaged PAC spectral densities for musicians and non-musicians during the 39 Hz entrainment condition (top left corner).** (a) Grand-averaged PAC values for each gamma and slow wave frequency combination in the spectrum. (b) Percentage cases with significant PAC (white; $p < 0.01$, and grey; $p < 0.05$) for each gamma and slow wave frequency combination in the spectrum. (c) and (d) Spectral densities for ‘musicians’ (red legend) and ‘non-musicians’ (blue legend).
